# A short perinuclear amphipathic α-helix in Apq12 promotes nuclear pore complex biogenesis

**DOI:** 10.1101/2021.08.13.456248

**Authors:** Wanlu Zhang, Azqa Khan, Jlenia Vitale, Annett Neuner, Kerstin Rink, Christian Lüchtenborg, Britta Brügger, Thomas H. Söllner, Elmar Schiebel

**Affiliations:** Zentrum für Molekulare Biologie der Universität Heidelberg, DKFZ-ZMBH Allianz, Im Neuenheimer Feld 282, 69120 Heidelberg, Germany; Biochemie-Zentrum der Universität Heidelberg, Im Neuenheimer Feld 328, 69120 Heidelberg, Germany; Cell Biology and Biophysics Unit, European Molecular Biology Laboratory (EMBL), Heidelberg, Germany

## Abstract

The integral membrane protein Apq12 is an important nuclear envelope (NE)/ER modulator that cooperates with the nuclear pore complex (NPC) biogenesis factors Brl1 and Brr6. How Apq12 executes these functions is unknown. Here we identified a short amphipathic α-helix (AαH) in Apq12 that links the two transmembrane domains in the perinuclear space and has liposome-binding properties. Cells expressing an *APQ12* (*apq12-ah*) version in which AαH is disrupted show NPC biogenesis and NE integrity defects, without impacting upon Apq12-ah topology or NE/ER localization. Overexpression of *APQ12* but not *apq12-ah* triggers striking over-proliferation of the outer nuclear membrane (ONM)/ER and promotes accumulation of phosphatidic acid (PA) at the NE. Apq12 and Apq12-ah both associate with NPC biogenesis intermediates and removal of AαH increases both Brl1 levels and the interaction between Brl1 and Brr6. We conclude that the short amphipathic α-helix of Apq12 regulates the function of Brl1 and Brr6 and promotes PA accumulation at the NE during NPC biogenesis.

## Introduction

The nuclear envelope (NE) is a double membrane consisting of the outer nuclear (ONM) and inner nuclear (INM) membranes that surround and protect the nucleus. The ONM is continuous with the endoplasmic reticulum (ER), contains attached ribosomes, carries attachment sites for cytoskeletal elements and shares components with the ER (reviewed in Hetzer et al., 2005). In addition, the ER and ONM are the sites of triacylglycerol (TAG) and steryl ester lipid biosynthesis (Sorger & Daum, 2003). These lipids are essential for membrane growth, and, in case of TAG, also for energy storage in form of cytoplasmic lipid droplets (Hashemi & Goodman, 2015, Thiam, Farese et al., 2013). The INM is involved in genome stability, chromatin organization and regulation of gene expression (Beck & Hurt, 2017, Onischenko, Tang et al., 2017, Ungricht & Kutay, 2017). A recent publication indicated that TAGs are also synthesised at the INM before being incorporated into nuclear lipid droplets (Romanauska & Kohler, 2018). Consistent with these distinct functions, the INM and ONM are specified by proteins and lipids that are district from one another.

The NPC is a large oligomeric complex containing about 30 different proteins that is embedded at sites at which the INM and ONM fuse. The NPC acts as a gateway at the partition between the cytoplasm with the nucleoplasm (Beck & Hurt, 2017). In human cells and other eukaryotes undergoing an open mitosis, NPCs assemble via two mechanistically distinct pathways. The post-mitotic pathway promotes the assembly of NPCs on the decondensing chromatin, shortly after anaphase onset (Otsuka, Steyer et al., 2018). During interphase, NPCs assemble by a distinct inside-out mechanism starting from within the nucleus at the INM, the so-called interphase pathway (Otsuka, Bui et al., 2016, Winey, Yarar et al., 1997, Zhang, Neuner et al., 2018). In budding yeast, with its closed mitosis, the interphase pathway is the only mechanism to assemble NPCs.

NPC biogenesis intermediates of the interphase pathway have been described in human cells by electron tomography (Otsuka et al., 2016). In wild type (WT) yeast cells, NPC intermediates have not been observed, probably because NPCs assembly is relatively fast and infrequent (∼2 assembly events per minute per cell (Winey et al., 1997)). However, mutations in genes coding for several nucleoporins (Nups) lead to the accumulation of, so called, herniations, deformations of the INM that are probably filled with Nups (Aitchison, Blobel et al., 1995, Murphy, Watkins et al., 1996, Rampello, Laudermilch et al., 2020, Wente & Blobel, 1993). Recently, we suggested that at least some of these herniations arise from defective parts of the NPC biogenesis pathways, for example the failure of the fusion of the INM with the ONM during the assembly process (Onischenko et al., 2017, Zhang et al., 2018).

Mechanistic principles of NPC complex biogenesis by the interphase pathway are poorly understood. In mammalian cells, early factors involved in interphase NPC biogenesis include Y-complex subunits NUP107 and NUP133, the INM protein SUN1 and the transmembrane nucleoporin POM121 (Doucet, Talamas et al., 2010, Otsuka & Ellenberg, 2018, Souquet, Freed et al., 2018, Talamas & Hetzer, 2011). In yeast, the two paralogous integral NE proteins Brl1 and Brr6 function in NPC biogenesis (de Bruyn Kops & Guthrie, 2001, Hodge, Choudhary et al., 2010, Lone, Atkinson et al., 2015, Saitoh, Ogawa et al., 2005, Tamm, Grallert et al., 2011, Zhang et al., 2018). Loss of function of Brl1 (<u>Br</u>r6 like protein No. <u>1</u>) or Brr6 (<u>b</u>ad response to refrigeration) gives rise to herniations without impacting upon the insertion or function of pre-existing NPCs to indicate that both proteins are specifically required for NPC assembly but not for the maintenance of assembled NPCs. Consistent with this notion, Brl1 and Brr6 only associate with NPC intermediates and not with fully assembled NPCs (Zhang et al., 2018). Brl1 and Brr6 physically and genetically interact with the small integral NE/ER protein Apq12 to reveal synergistic regulation by all three of these proteins (Hodge et al., 2010, Lone et al., 2015, Scarcelli, Hodge et al., 2007). Cells with a deletion in *APQ12* (*apq12Δ*) are cold sensitive for growth for unknown reason. Puzzlingly, at 37°C *apq12Δ* cells show a defect in NPC biogenesis despite relatively normal growth (Thaller, Allegretti et al., 2019).

A clear functional link between *BRL1* and NPC assembly has been revealed by genetic suppression of the physical interaction between the FG repeat containing Nup116 and the inner ring protein Nup188 that has a scaffolding function during NPC biogenesis leading to the accumulation of herniations in *NUP116* and *NUP188* defective cells (Allegretti, Zimmerli et al., 2020, Onischenko et al., 2017). Interestingly, the INM/ONM fusion defect of these scaffolding defective yeast cells was efficiently suppressed by overexpression of *BRL1* but not of *BRR6* (Zhang et al., 2018). This suggests a role of Brl1 in INM/ONM fusion and also shows that Brl1 and Brr6 have distinct functions during NPC biogenesis.

Here we report that Apq12 carries a short perinuclear amphipathic alpha-helix (AαH) connecting the two transmembrane domains. The AαH peptide binds to liposomes and amino acid substitutions that disrupt the amphipathic nature of the peptide abolish the binding to liposomes. Cells carrying an Apq12 with a defective AαH (*apq12-ah*) are cold sensitive for growth, show NPC biogenesis defects and disrupted NE even though the distribution of the mutant protein is unaffected and it assumes the correct topology within the membrane. Overexpression of *APQ12* triggers strong over-proliferation of the ONM and the accumulation of phosphatidic acid (PA) at the NE in a manner that is reliant upon the presence of a functional AαH. Apq12 associates with NPC biogenesis intermediates at bent INM segments, a localization that does not require a functional AαH. In addition, *apq12-ah* mutant shows elevated Brl1 and Brr6 levels and enhanced interaction between Brl1 and Brr6. Taken together, these data place the AαH of Apq12 into a strategic position to co- ordinate PA accumulation at the NE, Brl1-Brr6 interaction and NPC biogenesis.

## Results

### Apq12 carries a short lipid-binding AαH between the two transmembrane domains

Recently we have shown that the integral membrane protein Brl1 associates with NPC assembly intermediates and may promote fusion of the ONM with the INM during NPC biogenesis (Zhang et al., 2018). Brl1 interacts and cooperates with the integral membrane protein Apq12 (Lone et al., 2015). In order to gain an understanding of the molecular roles of Apq12 (Fig. 1A), we sought functional elements within this protein. The AmphipaSeeK program predicated an amphipathic α-helix (AαH) with two positively charged amino acids (Lys and Arg) on the hydrophilic side and hydrophobic amino acid residues on the opposite side of the helix (Fig. 1B; “Apq12”) (Sapay, Guermeur et al., 2006), between the two TM domains of Apq12 (Fig. 1A).

**Figure 1.**
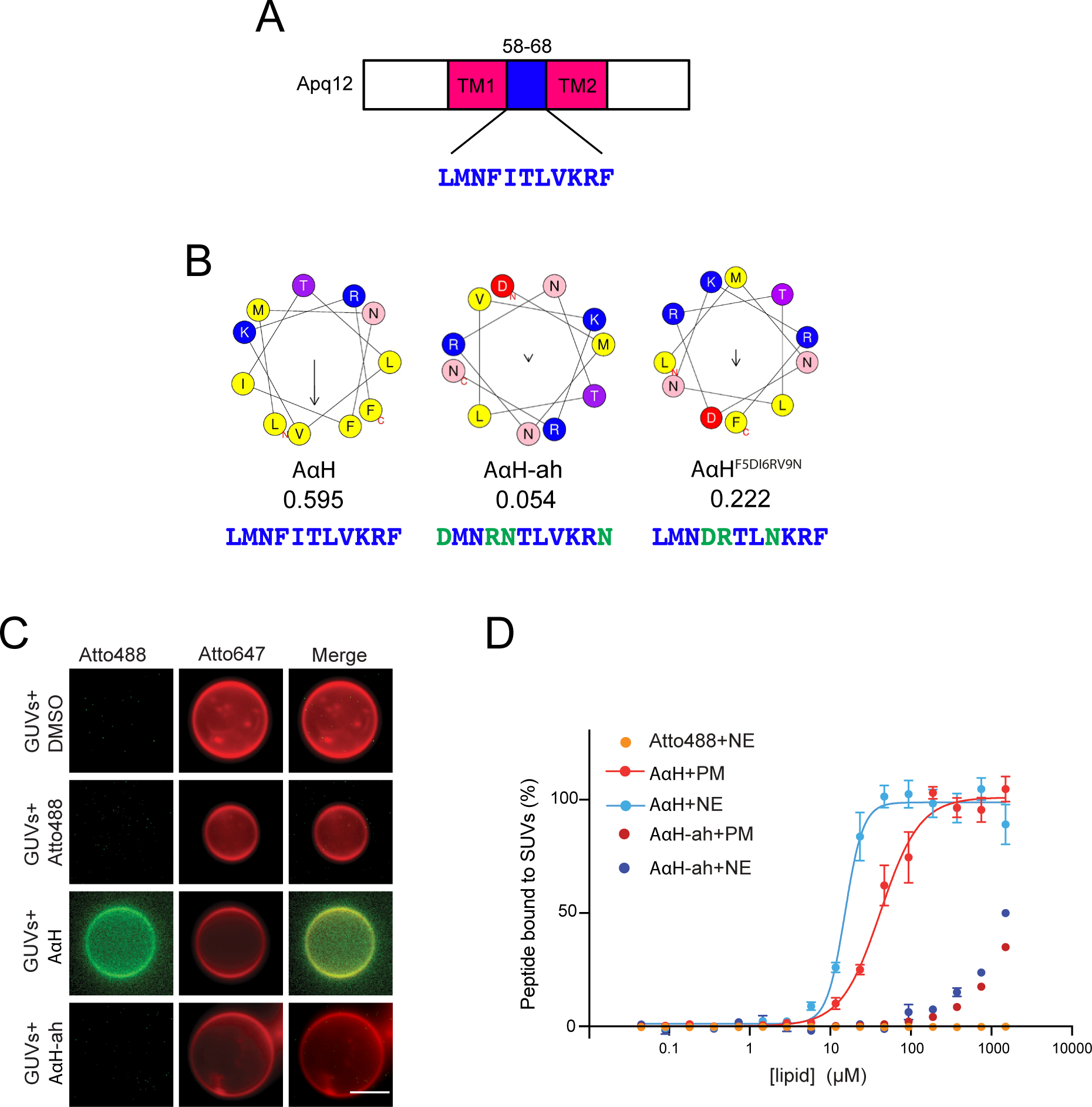
Apq12 contains an amphipathic helix in the luminal domain. **(A)** Domain organization of Apq12. The first transmembrane (TM1) domain, an AαH (blue) and the second transmembrane (TM2) domain are indicated. **(B)** Heliquest predictions of the AαH, AαH-ah, and AαH^F5DI6RV9N^ helices along with their hydrophobic moments. The amino acids marked in green indicate amino acid changes introduced in AαH in order to reduce the hydrophobic moment. **(C)** Binding of Atto488 labelled synthetic AαH and AαH-ah peptides to Atto647 labelled GUVs, *in vitro*. DMSO and Atto488 dye are used as controls. Scale bar: 5 µm. **(D)** Liposomes with different lipid compositions were titrated against Atto488 labeled AαH and AαH-ah peptides and the Atto488 dye as control. The data were normalized to the amount of bound peptide and the KD value of AαH peptide-liposome interaction was calculated from the Hill equation (PM: KD= 58.90 ± 5.35 µM, NE: KD= 15.90 ± 0.91 µM). The AαH-ah peptide and the Atto488 dye (together with NE lipids; same result was obtained with PM lipids) show no or only very weak MST signals. Values are given as means ± S.E.M, n = 3.

The positively charged amino acids of an AαH (Fig. 1B, marked blue) have the ability to interact with polar–apolar lipid surfaces while the hydrophobic interface (Fig. 1B, yellow) may interact with the aliphatic chains of the lipids (Gimenez-Andres, Copic et al., 2018). Consistent with this lipid binding prediction, we found that the Atto488 labelled synthetic Apq12 peptide (AαH) bound to uni-lamellar vesicles (GUVs; 1-10 µM in diameter; Fig. 1C). As control for the binding, we used the Atto488 dye alone, and introduced amino acid changes in the Atto488-labeled Apq12 peptide (AαH-ah) that abolished the helical hydrophobic moment, as an indication of the amphiphilicity of the helix (Eisenberg, Weiss et al., 1982), from 0.595 to 0.054 (Fig. 1B). The Atto488 dye and the AαH-ah peptide both failed to bind to liposomes (Fig. 1C).

To analyze the binding efficiency of AαH peptide in a more quantitative manner, SUV (small uni-lamellar vesicles; 80-120 nm, Fig. S1A) bound peptide was separated from the unbound peptide through a nycodenz gradient. Using SUVs composed of nuclear envelope (NE) and plasma membrane (PM) lipids, we tested the impact of the lipid composition. NE derived SUVs showed a higher AαH peptide binding efficiency compared to the PM SUVs (Fig. S1B).

The Atto488 dye and the AαH-ah peptide did not bind to the SUVs in this experimental regime (Fig. S1B). In addition, the peptide without liposomes was unable to float through the nycodenz gradient (Fig. S1B, control).

Next, SUV binding of the AαH peptide was quantified using microscale thermophoresis (MST) measurements. The KD of AαH peptide binding to NE-lipid derived SUVs was 16 µM (Fig. 1D). The binding affinity of the AαH peptide to PM derived SUVs was decreased to KD = 59 µM (Fig. 1D). Importantly, AαH-ah peptide failed to bind to any of these SUVs with a measurable KD. Thus, the AαH peptide binds to liposomes depending on its amphipathic nature and the lipid composition of the liposomes.

### The AαH of Apq12 localizes in the perinuclear space and is not required for subcellular localization and topology of Apq12

We analyzed whether the amphipathic nature of the AαH is important for the function of Apq12. Since *apq12Δ* mutants show a growth defect at 16°C (Scarcelli et al., 2007), we tested *APQ12*, *apq12Δ, apq12-ah* and an additional AαH *APQ12* mutant*, apq12^F5DI6RV9N^*, for growth at different temperatures. The helical hydrophobic moment of the AαH was decreased in AαH^F5DI6RV9N^ to 0.222 and therefore has an intermediate value between the WT AαH of Apq12 and AαH-ah of Apq12-ah (Fig. 1B). *apq12-ah* mutant completely failed to grow at 16°C and showed reduced growth at 23°C, similar to the *apq12Δ* cells (Fig. 2A). In contrast, growth of *apq12^F5DI6RV9N^* mutant cells was only reduced, but not completely abolished, at 16°C (Fig. 2A). Therefore, all further experiments were performed with *apq12- ah* cells. Thus, the amphipathic nature of the AαH in Apq12 is important for cell growth at lower temperatures.

**Figure 2.**
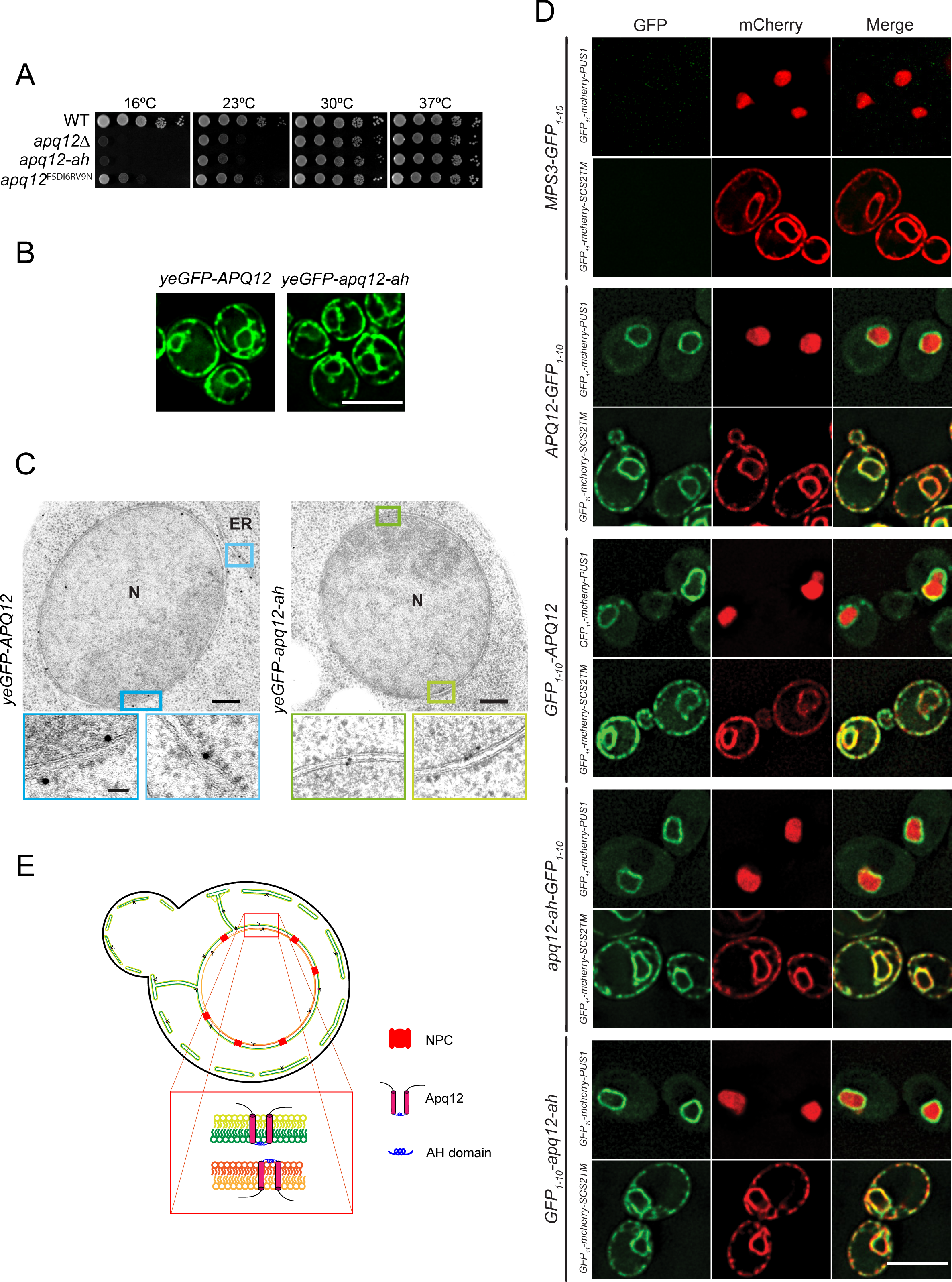
The AαH of Apq12 is located in the perinuclear space and does not influence the subcellular localization and topology. **(A)** Growth test of WT *APQ12*, *apq12Δ*, *apq12-ah, apq12*^F5DI6RV9N^ mutants at the indicated temperatures. 10-fold serial dilutions were spotted onto YPAD plates. **(B)** Localization of yeGFP-Apq12 and yeGFP-apq12-ah were analyzed by fluorescence microscopy. Scale bar: 5 µm. **(C)** yeGFP-Apq12 and yeGFP-apq12-ah localization by immuno-EM. Gold particles (10 nm) indicate the localization of yeGFP-Apq12 and yeGFP-apq12-ah at the NE and ER. The rectangles indicate the enlargements that are shown underneath. Abbreviation: N: nucleus. Scale bars: 200 nm and enlargements 50 nm. **(D)** Strains carrying C- and N- terminal fusions of Apq12 and Apq12-ah with GFP1-10 were imaged to check for reconstitution of GFP with GFP11 from GFP11-mCherry-Scs2TM (ER reporter; GFP11 in the cytoplasm) and GFP11-mCherry-PUS1 (nuclear reporter). Mps3-GFP1-10 is used as a negative control. Scale bar: 5 µm. (**E**) Localization and topology model for Apq12.

We next analyzed whether the subcellular localization of Apq12 requires the AαH. Consistent with published data on Apq12 distribution (Lone et al., 2015), yeGFP-Apq12 localized along the NE and the cell periphery, and a location that is probably the peripheral ER (Fig. 2B). Similar localizations were observed for yeGFP-Apq12-ah (Fig. 2B). Thus, the integrity of the AαH of Apq12 is not important for the subcellular distribution of the protein.

Immuno-electron microscopy of yeGFP-Apq12 and yeGFP-Apq12-ah using GFP antibodies and protein A-gold detected both proteins along the INM, ONM, cytoplasmic and cortical ER (Fig. 2C and Fig. S1C). Apq12 was detected with the same frequency at the INM as at the ONM (Fig. S1C). For Apq12-ah there was a mild enrichment at the INM over the ONM (Fig. S1C). In addition, yeGFP-Apq12 and yeGFP-Apq12-ah associated with 10% and 16% of NPCs, respectively, while most NPCs were not labelled (Fig. S1D). This may indicate a transient association of the proteins with assembling NPCs, as is the case for Brl1 and Brr6 (Zhang et al., 2018). In summary, Apq12 and Apq12-ah show similar subcellular localizations.

Apq12 is a protein of the NE and the ER with two predicted membrane spanning regions (TM1 and TM2) that are connected by the AαH (Fig. 1A). To determine the topology of Apq12 and whether it requires a functional AαH, we measured the accessibility of the N- and C-terminus of Apq12 and Apq12-ah by two approaches. First, we used the split GFP system for assessment of the localization of the N- and C-termini of Apq12 (Smoyer, Katta et al., 2016). The overlapping localization of GFP11 and GFP1-10 restores GFP fluorescence. Thus, by combining GFP1-10 tagged versions of a protein with nuclear and cytoplasmic GFP11 localised proteins, we can determine the topology of a protein. As a control, we used the perinuclear space localization of the C-terminus of the SUN-domain protein Mps3-GFP1-10 that failed to restore a fluorescent GFP signal when co-expressed with the ER and ONM localized GFP11-mCherry-Scs2TM that carries GFP11 at the ONM/ER on the cytoplasmic side and the nuclear GFP11-mCherry Pus1 (Fig. 2D, top panel) consistent with published data (Smoyer et al., 2016). Interestingly, Apq12-GFP1-10 and GFP1-10-Apq12 combined with GFP11-mCherry Pus1 resulted in a green fluorescent NE signal and in case of GFP11- mCherry-Scs2TM a NE/ER signal. These data show that the N- and C-termini of Apq12 are localized to either the cytoplasm or the nucleoplasm depending on whether Apq12 is at the ONM/ER or INM, respectively (Fig. 2D, middle). Very similar results were obtained for Apq12-ah (Fig. 2D, bottom).

As a further probe of Apq12 topology, we took advantage of the fact that biotin ligases are not contained in the perinuclear space, and assessed the ability of the N- or the C-termini of Apq12 and Apq12-ah to be biotinylated when fused with the histidine-biotin-histidine (HBH) tag as a topology marker (Zhang et al., 2018). Pom152 that contains a short cytosolic N-terminal region, one TM domain and a C-terminal region in the perinuclear space was used as a control (Tcheperegine, Marelli et al., 1999). Consistent with the topology of Pom152, only the N-terminal HBH tag but not the C-terminal tag of Pom152 was biotinylated, demonstrating that the HBH biotinylation approach identifies the topology of NE proteins correctly (Fig. S1E). Both the N- and C-termini of Apq12 and Apq12-ah were biotinylated when fused to the HBH tag that carries a biotin acceptor site (Fig. S1E), confirming the findings from the split GFP approach.

In conclusion, since N- and C-termini of Apq12 localized to either the cytoplasm or nucleoplasm, the AαH resides in the perinuclear space where it connects the two TM domains (Fig. 2E). A functional AαH is not required for this topological arrangement of Apq12.

### The AαH of Apq12 is important for NE integrity, NPC biogenesis and lipid homeostasis

In order to understand how loss or impairment of the Apq12 function affects the NE and NPCs, we analyzed the phenotypes of *APQ12*, *apq12Δ* and *apq12-ah* cells by electron microscopy (EM). As expected, *APQ12* wild type cells had spherical nuclei with intact NE (Fig. 3A). Herniations as an indication of a defect in NPC biogenesis were detected as major phenotype in *apq12Δ* (Fig. 3B and C, nucleus marked by red square) and *apq12-ah* mutants at 37°C (Fig. 3D and E, nucleus marked by red square). About 70% of the *apq12Δ* mutants incubated at 23°C showed invaginations of the NE (Fig. 3C). Herniations were relatively infrequent at this growth temperature. Invaginations of the NE and NE breakdown were the major defects of the *apq12-ah* mutant at 23°C (Fig. 3E). At 16°C, the major defects of *apq12Δ* cells were NE invaginations and NE breakdown followed by herniations (Fig. 3B and C). *apq12-ah* mutant showed NE breakdown and extrusions as major defects at 16°C (Fig. 3D and E). Taken together, *apq12Δ* and *apq12-ah* mutants show phenotypic variations indicating that inactivation of the AαH does not cause the complete loss of Apq12 function. In addition, the consequence of Apq12 or AαH loss depends on the temperature. The rupture of the NE explains the lethality of *apq12Δ* and *apq12-ah* cells at 16°C. Loss of AαH function triggers a defect in NPC biogenesis at 37°C.

**Figure 3.**
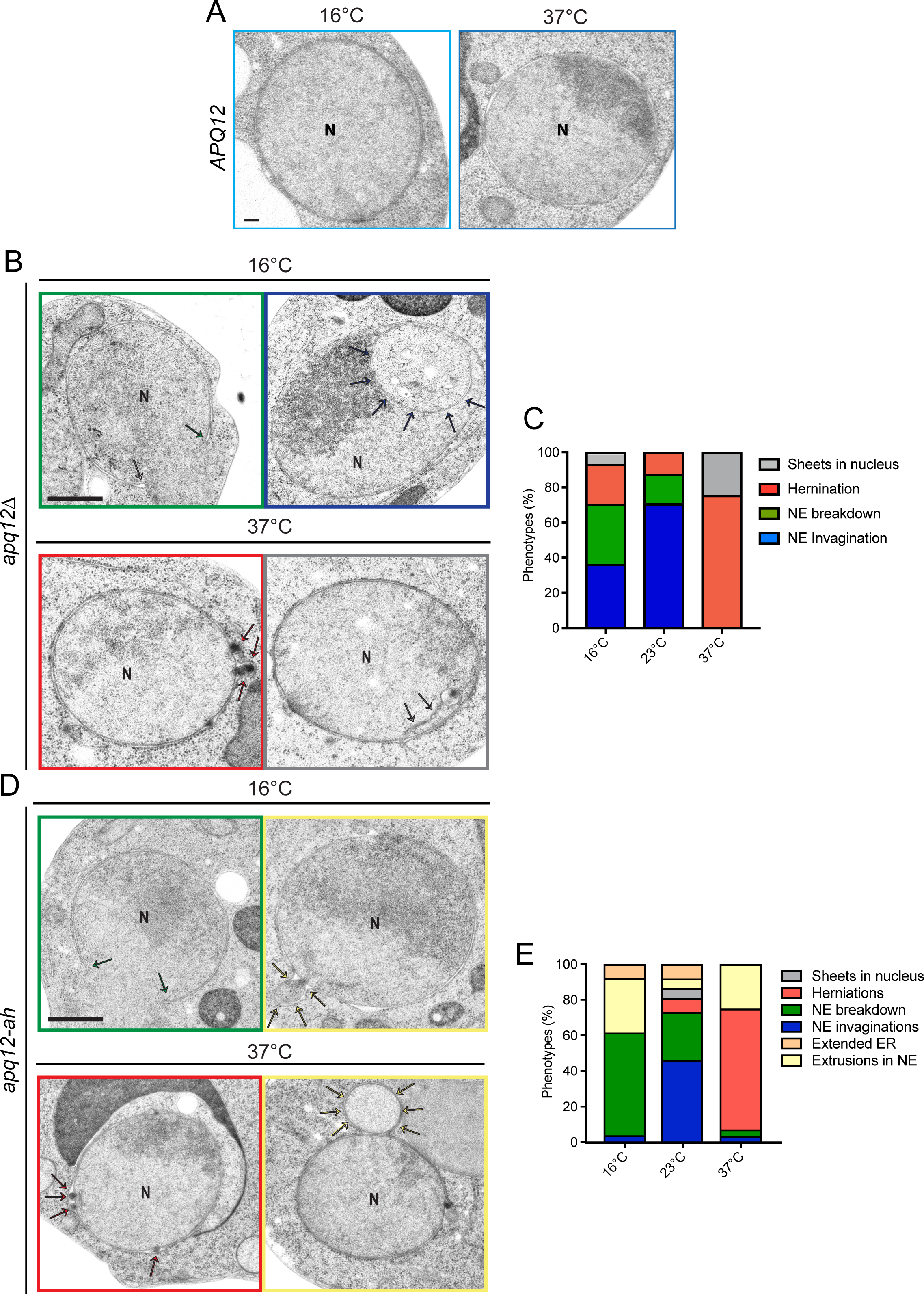
Phenotypes of *apq12Δ* and *apq12-ah* cells. (A) EM analysis of *APQ12* WT cells incubated at the indicated temperatures. (B) EM analysis of *apq12Δ* mutants incubated at the indicated temperatures. The arrows point towards the phenotype that is indicated by the colour code encircling the picture. (C) Quantification of phenotypes from (B). Cells (n=44/16°C, 40/23°C and 57/37°C) were analyzed per temperature. (D) EM analysis of *apq12-ah* mutants grown at the indicated temperature. The arrows point towards the phenotype that is indicated by the colour code encircling the picture. (E) Quantification of phenotypes from (D). Cells (n=27/16°C, 39/23°C and 28/37°C) were analyzed per temperature. (A, B and D) Abbreviation: N, nucleus. Scale bars: (A) 100 nm; (B, D) 500 nm.

It has been suggested that Apq12 has a function in lipid homeostasis (Lone et al., 2015), which might then account for the NE and NPC defects of *apq12* and *apq12-ah* mutants. Therefore, we directly asked whether the AαH of Apq12 has an impact on lipid homeostasis, by incubating WT, *apq12Δ* and *apq12-ah* cells at 16°C, 30°C and 37°C and assessing their cellular lipid content by mass spectrometry. Overall *apq12Δ* and *apq12-ah* mutants showed comparable lipid changes relative to WT (Fig. S2). The increase in ergosteryl ester (EE) and TAG and a decrease in ergosterol (Erg) and most species of glycerophospholipids (GPL) were prominent phenotypes in *apq12Δ* and *apq12-ah* mutants (Fig. S2A and B), indicating a lipid metabolism flow from membrane lipids to storage lipids. In addition, we observed a significant decrease of double bonds in GPL in *apq12Δ* and *apq12-ah* mutants (Fig. S2C). Moreover, the chain length (>34) in GPL was significantly increased in both *APQ12* mutants (Fig. S2D). The reduction of membrane lipids, the decrease in the number of double bonds and the increase of chain length in GPL indicate a decrease in membrane fluidity in *apq12Δ* and *apq12-ah* mutants, explaining the defects in NE breakdown at 16°C when the flexibility of membrane lipids will be reduced by the reduction in kinetic energy. These data suggests that the AαH of Apq12 does play a role in maintaining lipid homeostasis.

### Increased Apq12 levels are toxic to cells and cause the mis-localization of the NPC biogenesis factors Brl1 and Brr6 and the ER proteins Sec63 and Ole1

To gain deeper insights into the function of Apq12, we overexpressed *APQ12* using the galactose-inducible PGal1 promoter and followed the growth of these modified yeast cells. Because we lack an Apq12 antibody, we tagged Apq12 with 6His (Apq12-6His) to support immuno-detection of the fusion protein. Overexpression of *APQ12* and *APQ12-6His* was equally toxic for cells (Fig. 4A). Such overexpression toxicity was not observed for the partner proteins *BRL1* and *BRR6* which also encode integral membrane proteins (Fig. 4B). Thus, of the three proteins in this functional module, only *APQ12* has toxic consequences upon overproduction.

**Figure 4.**
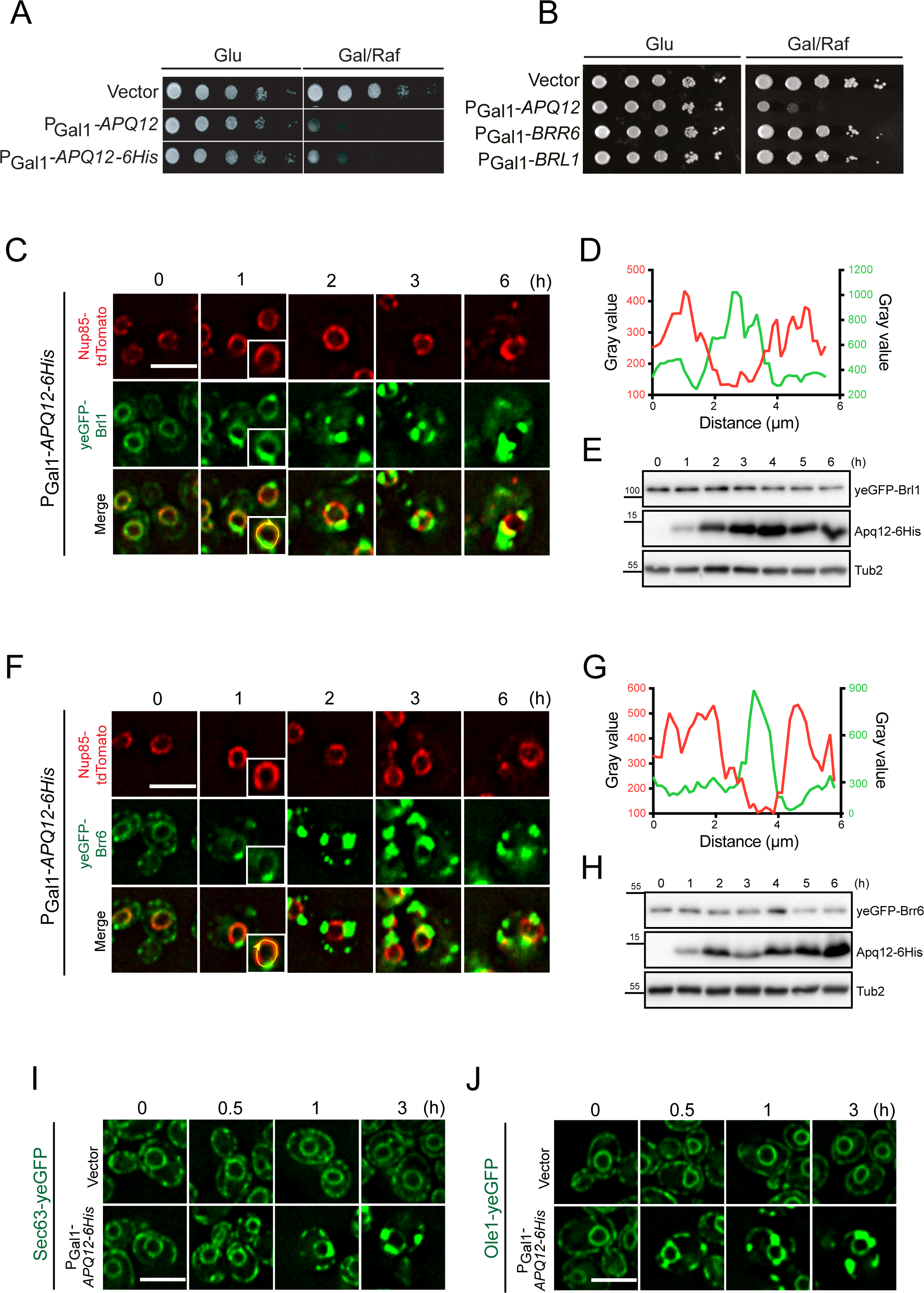
Overexpression of *APQ12* causes the mislocalization of ER proteins. (A) Overexpression of *APQ12* is toxic for the cells. WT cells with the vector control, PGal1-*APQ12* or PGal1-*APQ12-6His* were spotted in 10-fold serial dilutions onto YPAD (Glu) and YARaf/Gal (Gal/Raf) plates at 30°C. (B) Of the *APQ12*, *BRL1*, *BRR6* module only *APQ12* overexpression is toxic. WT cells with the vector control, PGal1- *APQ12*, PGal1-*BRR6* and PGal1-*BRL1* were spotted in 10-fold serial dilutions onto YPAD (Glu) and YARaf/Gal (Gal/Raf) plates at 30°C. (C) Overexpression of *APQ12* causes mislocalization of yeGFP-Brl1. Cells with either the vector control (Fig. S3C) or PGal1-*APQ12-6His* were incubated with galactose for the indicated times. The boxed cell at 1 h is a two-fold enlargement of the selected cell. (D) Line scan along the NE of a PGal1-*APQ12-6His yeGFP-BRL1 NUP85-tdTomato* cell (enlarged boxed cell in (C)) incubated for 1 h with galactose. It shows that yeGFP-Brl1 and Nup85- tdTomato are localized on separate domains along the NE. (E) Immunoblot of PGal1- *APQ12-6His yeGFP-BRL1 NUP85-tdTomato* cells. The pGal1 promoter was induced by the addition of galactose (t=0). Samples were taken after the indicated times. Tub2 is a loading control. Apq12-6His was detected by anti-His antibodies. (F) Overexpression of *APQ12* causes mislocalization of yeGFP-Brr6 and Nup85- tdTomato. Cells with either the vector control (Fig. S3D) or PGal1-*APQ12-6His* were incubated with galactose for the indicated times. The boxed cell at 1 h is a two-fold enlargement of the selected cell. (G) Line scan along the NE of a PGal1-*APQ12-6His yeGFP-BRR6 NUP85-tdTomato* cell (enlarged cell in (F)) incubated for 1 h with galactose. (H) Immunoblot of PGal1-*APQ12-6His yeGFP-BRR6 NUP85-tdTomato* cells. The pGal1 promoter was induced by the addition of galactose (t=0). Samples were taken after the indicated times. Tub2 is a loading control. (I, J) Overexpression of *APQ12* causes mislocalization of the ER protein Sec63-yeGFP (I) and Ole1- yeGFP (J). Cells with either the vector control or PGal1-*APQ12-6His* were incubated with galactose for the indicated time. (C, F, I and J) Scale bars: 5 µm.

We next tested whether overexpression of *APQ12* and *APQ12-6His* impact on NE structure and function. We started by analyzing the impact of *APQ12* over-expression on the distribution of the NPC biogenesis factors Brl1 and Brr6 (Fig. 4C-H; Fig. S3A and B). In vector control cells, yeGFP-Brl1 and yeGFP-Brr6 exhibited a uniform distribution throughout the NE (Fig. S3C and D). In contrast, one hour of induction of the PGal1 promoter to elevate Apq12 levels led to dense clustering of Brl1 and Brr6 on the NE (Fig. 4C-H, Fig. S3A and B). These clusters were devoid of the NPC marker Nup85-tdTomato (Fig. 4C, D, F and G, Fig. S3A and B). Clustering of yeGFP-Brl1 and yeGFP-Brr6 in response to *APQ12-6His* overexpression was also seen in time-lapse experiments (Fig. S3E and F). Because PGal1- *APQ12* and PGal1-*APQ12-6His* caused similar defects, we used PGal1-*APQ12-6His* in further experiments.

We next asked how *APQ12-6His* overexpression affected the distribution of proteins at the NE/ER and INM, using the NE/ER proteins Sec63 and Ole1, the INM protein Asi3 and the ER luminal marker dsRED-HDEL as exemplars to test the impact upon NE/ER proteins (Feldheim, Rothblatt et al., 1992, Khmelinskii, Blaszczak et al., 2014). As seen for yeGFP- Brl1 and yeGFP-Brr6, the smooth distribution of Sec63-yeGFP and Ole1-yeGFP along the NE was transformed into clusters as early as 1 h of PGal1-*APQ12-6His* induction (Fig. 4C, F, I and J). The localization of INM protein Asi3-yeGFP was initially unaffected by *APQ12* overexpression, at the 1 h time point, however, by 3 h of PGal1 induction, modest clustering of Asi3-yeGFP along the NE emerged (Fig. S4A; white arrows). The ER marker dsRED-HDEL, that in vector control cells uniformly stained the NE and the cortical ER, started to accumulate in clusters within an hour of PGal1-*APQ12-6His* induction (Fig. S4B). Thus, increased Apq12 levels appears to impact upon the subcellular localization of ONM/ER proteins more strongly than the INM protein Asi3.

### The luminal AαH of Apq12 contributes to toxicity

We asked whether AαH is important for the deformation of the NE. PGal1-*apq12-ah-6His* overexpression was less toxic than PGal1-*APQ12-6His* (Fig. 5A) even though both proteins were overexpressed to similar levels (Fig. S5A). Consistent with the strongly reduced toxicity of PGal1-*apq12-ah-6His*, yeGFP-Brl1 and yeGFP-Brr6 clusters formed later and were notably less prominent in PGal1-*apq12-ah-6His* cells compared to PGal1-*APQ12-6His* cells (Fig. 5B-E). Thus, an intact AαH is a major factor in the toxic impact of *APQ12* overexpression.

**Figure 5.**
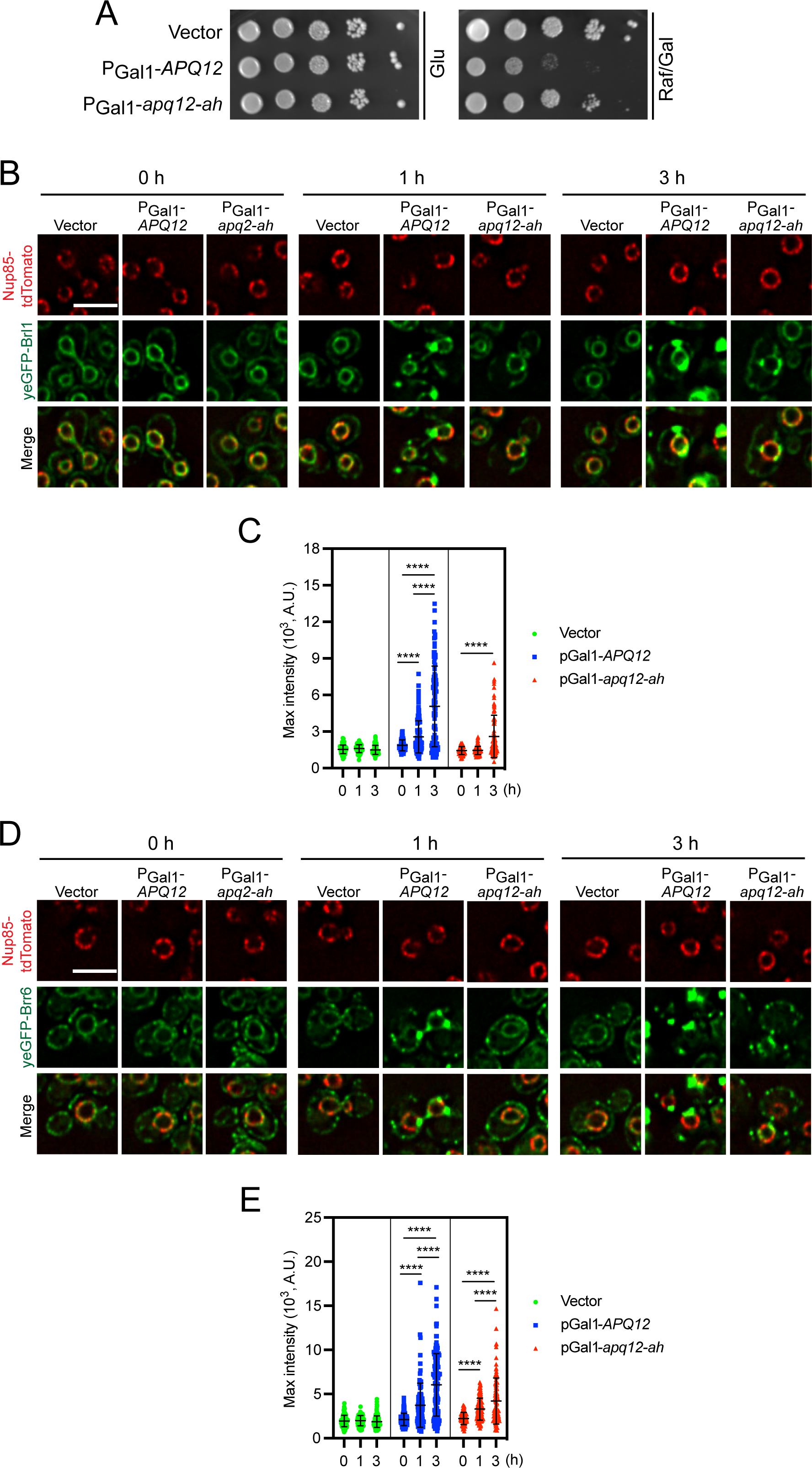
The amphipathic helix of Apq12 is important for membrane remodelling. (A) Inactivation of the amphipathic α-helix of Apq12 reduces overexpression toxicity. Growth test of cells containing PGal1-*apq12-ah*, in comparison to PGal1-*APQ12* and vector control cells. 10-fold serial dilutions of cells were spotted onto glucose (Glu) and galactose/raffinose (Gal/Raf) plates at 30°C. (B-E) Impact of *APQ12* and *apq12-ah* overexpression on (B, C) *yeGFP-BRL1 NUP85-tdTomato* and (D, E) *yeGFP-BRR6 NUP85-tdTomato* cells. (C, E) are quantifications of the maximum GFP intensity of individual cells from (B) and (D) over time. Two independent experiments were performed, and more than 60 cells were analyzed per experiment for each time point. Values are given as means +/- SD. t-test ****p<0.0001. (B, D) Scale bars: 5 µm.

### Apq12 needs a functional AαH for the over-proliferation of the ONM and ER

To understand how *APQ12* overexpression leads to the changes on the ONM and ER, we performed EM analysis of PGal1-*APQ12-6His* and PGal1-*apq12-ah-6His* cells. Before the addition of galactose, the NE had uniform spherical morphology (Fig. S5B). Induction of PGal1-*APQ12-6His* led to a shift in the number of NE extensions (Fig. 6A-a, arrow) that were, in some cases, connected to the cortical ER (Fig. 6A-b) from 5% to 40% (Fig. 6B). After 1 and 3 h of galactose addition, PGal1-*APQ12-6His* cells contained ONM encircled vesicles containing granular material (Fig. 6A-c-f, red asterisks, and Fig. 6B). In contrast abnormal morphologies were not detected at the INM until 3 h after the induction of *APQ12* overexpression, whereupon the INM formed small buds into the luminal space of the ONM vesicles (Fig. 6A-e, arrow). The consequences of *apq12-ah-6His* overexpression were less severe in comparison to *APQ12-6His*. PGal1-*apq12-ah-6His* induction for 0.5 h and 1 h had no visible impact on the ONM or the ER in most cells (Fig. 6A-a’-d’). Only 3 h of PGal1-*apq12-ah- 6His* induction caused proliferation of the ONM in ∼25% of the cell sections (Fig. 6A-e’, arrows and Fig. 6B). This delayed accumulation of phenotypes in *apq12-ah-6His* cells is consistent with the reduced clustering of the *BRL1-yeGFP* and *BRR6-yeGFP* and impact on growth compared to *APQ12-6His* (Fig. 5).

**Figure 6.**
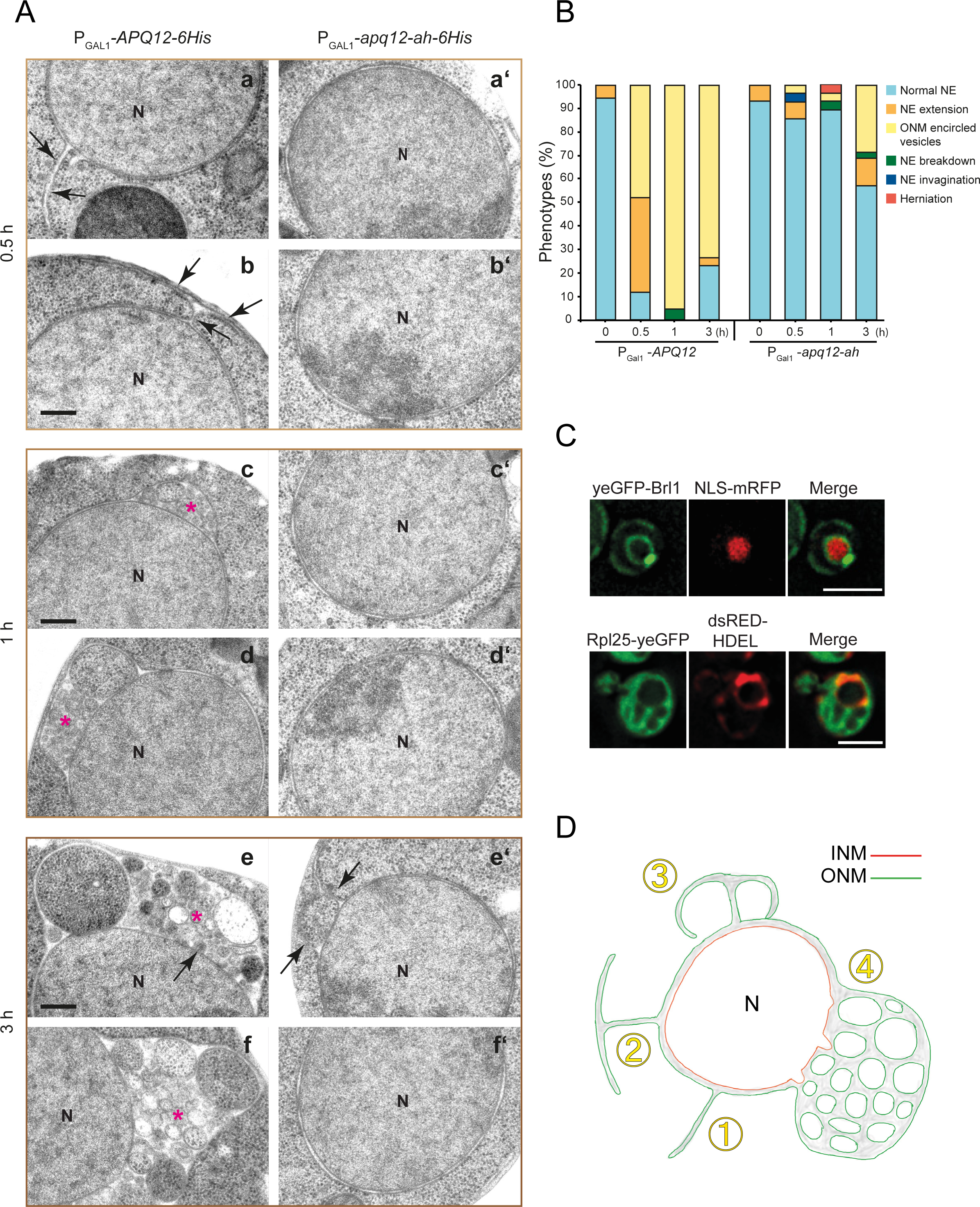
Overexpression of *APQ12* triggers hyper-proliferation of the ONM and ER. (A) EM analysis of WT cells expressing PGal-*APQ12-6His* (left) or PGal-*apq12-ah-6His* (right) by the addition of galactose for 0.5 h, 1 h, and 3 h. Arrows are explained in the result section; red asterisks indicate ONMs encircled vesicles containing granular material. Abbreviation: N, nucleus. Scale bar: 200 nm. (B) Quantification of phenotypes from (A). n = 25 cells were analyzed for each time point and strain. (C) PGal1-*APQ12* was overexpressed for 3 h by the addition of galactose. Top panel: cells expressing yeGFP-Brl1 along with a nuclear marker NLS-mRFP. Bottom panel: cells expressing yeGFP tagged ribosomal subunit Rpl25, and an ER luminal marker dsRED-HDEL. Scale bar: 3 µm (D) Model for the formation of membrane structures in response to *APQ12* overexpression. See result for details.

Overexpression of *APQ12* causes the accumulation of large, granular ONM encircled vesicles. To investigate the origin of the content in these vesicles, we expressed PGal1- *APQ12-6His* in cells with yeGFP tagged Brl1 as marker for the vesicles, and NLS-mRFP as nuclear marker. Overexpression of Apq12-6His induced yeGFP-Brl1 areas that were devoid of NLS-mRFP (Fig. 6C, top) indicating that these vesicles did not contain nuclear components. In a complementing experiment, we expressed PGal1-*APQ12-6His* in *RPL25- yeGFP dsRed-HDEL* cells with the green fluorescent ribosomal subunit as cytoplasmic marker and dsRed-HDEL as ONM/ER marker (Fig. 6C, bottom). Indeed, the dsRed-HDEL ONM extrusions co-localized with Rpl25-yeGFP, demonstrating that these vesicles contained cytoplasmic components.

In a time-lapse experiment, we analyzed the cellular distribution of Apq12-yeGFP and Apq12-ah-yeGFP upon PGal1 expression (Fig. S5C). Apq12-yeGFP accumulated most prominently at the NE extensions (Fig. S5C, 30 min, arrowheads). After 40 min of induction Apq12-yeGFP localized in larger spots (Fig. S5C, arrowheads), consistent with the accumulation of filled ONM deformations seen by EM (Fig. 6A). For the first 40 min of PGal1 induction Apq12-ah-yeGFP showed comparable localizations to Apq12-yeGFP (Fig. S5C). However, due to its defective AαH, Apq12-ah-yeGFP did not accumulate into larger spots. Analysis of PGal1-*APQ12-yeGFP* after 0.5 and 3 h of galactose addition by immuno-EM detected Apq12-yeGFP locating at NE extensions and ONM enriched vesicles (Fig. S5D). Thus, the enrichment of Apq12 at ONM/ER sites induces membrane proliferation.

When taken together, *APQ12* overexpression promotes extension of ER tubes from the ONM (Fig. 6D; steps 1 and 2). Fusion of these extensions with the NE (step 3) may entrap cytoplasmic content into ONM encircled compartments (step 4).

### *APQ12* triggers PA accumulation at the NE dependent on its AαH

The experiments above indicate that Apq12 promotes over-proliferation of the ONM and the ER in an AαH dependent manner. A mobilization of lipids by Apq12 would explain the ONM/ER expansion. To test for this possibility, we applied lipid mass spectrometry analysis to evaluate how PGal1-induced overexpression of *APQ12* and *apq12-ah* affected the composition of cellular lipids. Cells carrying the pGal1 construct were used as vector control. Samples were analyzed at 0, 1 and 3 h of galactose addition (Fig. S6A-D). Throughout the experiment the vector control behaved in a similar manner to *apq12-ah* cells. Interestingly, PGal1-*APQ12-6His* overexpression triggered an increase in DAG and TAG after 1 h of promoter induction (Fig. S6A). PS and EE in the PGal1-*APQ12-6His* sample was increased after 3 h induction. In addition, the number of two double bonds in glycerophospholipids (GPL) decreased in the PGal1-*APQ12-6His* sample after 1 h but with an increase of one double bond after 3 h. Furthermore, PGal1-*APQ12-6His* expression also affected the chain length of GPL after 3 h of induction compared to PGal1-*apq12-ah-6His* induction (Fig. S6D). Thus, PGal1-*APQ12-6His* expression has a mild impact on the lipid composition of cells.

Lipid mass spectrometry analysis did not enable us to make conclusions about local changes in lipid content, for example understanding whether the changes were restricted to one location such as the NE or cytoplasmic membrane systems. Using recently reported lipid sensors (Romanauska & Kohler, 2018), we tested whether *APQ12* overexpression affects PA accumulation at the INM or ONM and compared the outcome of overproduction of the wild type molecule with the impact of *apq12-ah* overexpression. We used the cytoplasmic Q2-mCherry and nuclear PA sensor NLS-Q2-mCherry for the analysis of PA changes at the ONM and INM, respectively. Consistent with published data (Romanauska & Kohler, 2018), at t=0 strong Q2-mCherry decoration of the cytoplasmic membrane (Fig. 7A and B) was accompanied by a weak Q2-mCherry nuclear signal. After 1 h and 3 h of PGal1-*APQ12* induction, the Q2-mCherry signal accumulated in a rim-like pattern at the NE and co- localized with developing yeGFP-Brl1 clusters at the NE (Fig. 7A-C). In PGal1-*apq12-ah* cells at t=0 the Q2-mCherry reporter showed similar localization to PGal1-*APQ12* cells (Fig. 7A) and induction for 1 h did not impact the localization of the Q2-mCherry sensor. After 3 h of induction the nuclear Q2-mCherry signal was nearly completely diminished without directing the sensor to the NE (Fig. 7A-C).

**Figure 7.**
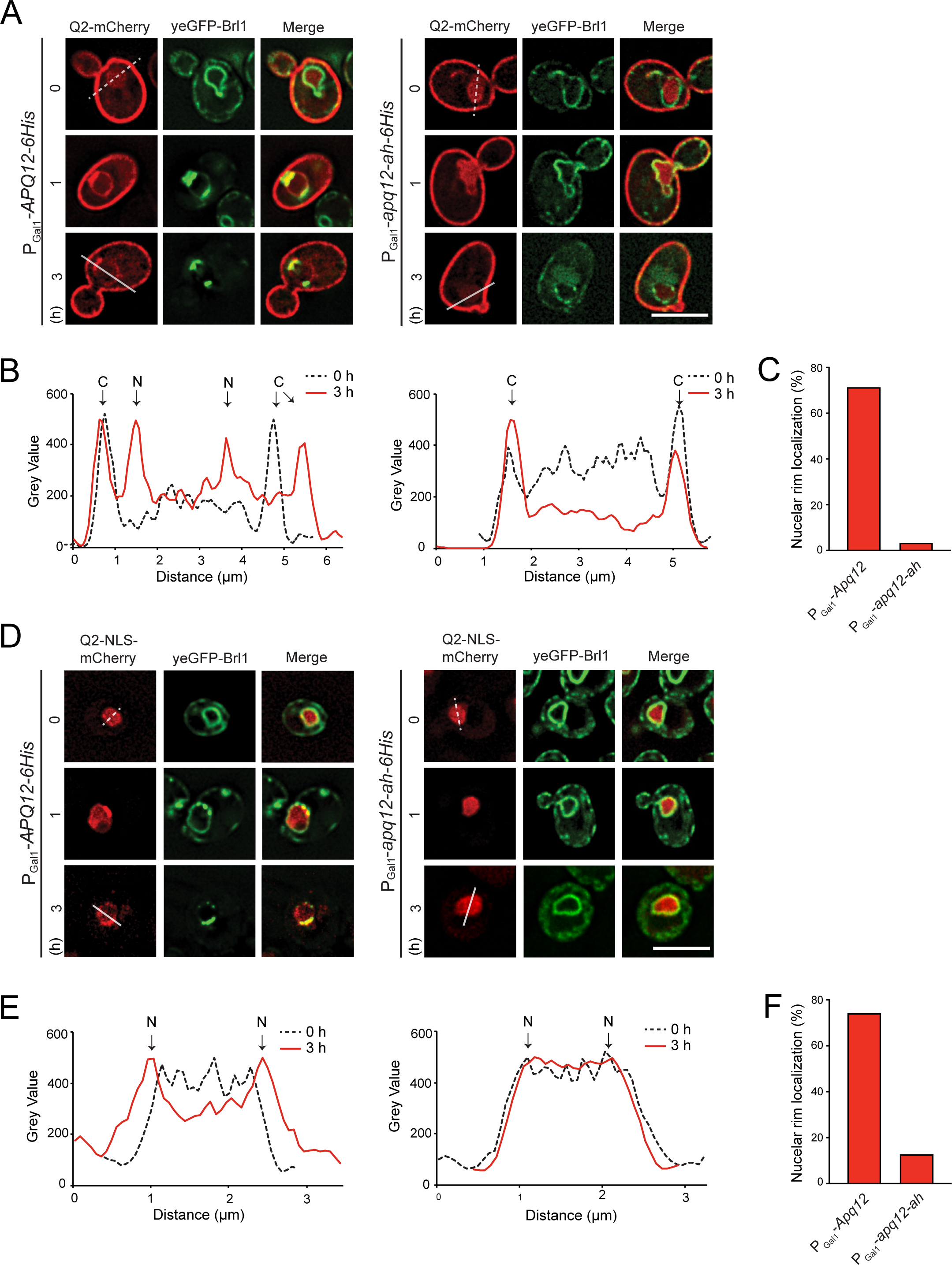
*APQ12* overexpression induces AαH dependent PA accumulation at the NE. (A) Localization of the cytoplasmic PA sensors (Q2-mCherry) in response to PGal1-*APQ12* and PGal1-*apq12-ah* induction at 0, 1 and 3 h. (B) Line scans of mCherry signal along the indicated lines across the cells in (A). Arrowheads mark the peaks corresponding to the cytoplasmic membrane (C) and NE (N). (C) Quantification of data from (A) showing percentage of cells with nuclear rim localization after 3 h of PGal1-*APQ12* induction. 100 cells carrying either PGal1-*APQ12* or PGal1-*apq12-ah* from one representative experiment were analyzed for nuclear rim localization of the PA sensor. (D) Localization of the nuclear PA sensors (NLS-Q2-mCherry) in response to PGal1-*APQ12* (left) and PGal1-*apq12-ah* (right) induction at 0, 1 and 3 h. (E) Line scans of mCherry signal along the indicated lines across the nuclei in (D). Arrowheads mark the peaks corresponding to the NE (N). (F) Quantification of data from (D) showing percentage of cells with nuclear rim localization done as in (C). (A, D) Scale bars: 5 µm.

Most interestingly, the nuclear NLS-Q2-mCherry (t=0) was recruited from a nuclear localization at t = 0 to a rim-like distribution at the NE 1-3 h after induction of PGal1-*APQ12- 6His* (Fig. 7D-F). In addition, NLS-Q2-mCherry strongly co-localized with yeGFP-Brl1 enrichments at the NE (Fig. 7D). In contrast, PGal1-*apq12-ah* expression did not impact the nuclear localization of NLS-Q2-mCherry (Fig. 7D-F). Thus, the AαH of *APQ12* is required to induce PA enrichment at the NE.

### The AαH of Apq12 is not required for the association of the protein with herniations

The co-localization of Apq12 and apq12-ah with a small subset of NPCs (Fig. S1D) raises the possibility that both proteins resemble Brl1 and Brr6 in associating with the newly assembled NPCs but not fully assembled NPCs (Zhang et al., 2018). We tested this notion by studying Apq12 and Apq12-ah localization in cells carrying temperature dependent *td-brl1* and *td-brr6* degrons, that accumulate NPC biogenesis intermediates at 37°C (Zhang et al., 2018). Localization of Apq12 and Apq12-ah at herniations in *nup116* cells was not analyzed because of the synthetically lethal phenotype of *nup116 apq12* mutant combination (Scarcelli et al., 2007). Interestingly, in *td-brl1* and *td-brr6* cells incubated at 37°C, yeGFP-Apq12 co- localized with Nup85-tdTomato while this was not the case in the equally treated WT control cells, as indicated by the scan of the NE signal (Fig. 8A and B). Quantification of the co - localization of yeGFP-Apq12 and Nup85-tdTomato signals along the NE showed a clear correlation between the Nup85-tdTomato and yeGFP-Apq12 signals in *td-brr6* and *td-brl1* cells, but not in WT control cells (Fig. 8C). We next asked whether a functional AαH is required for the co-localization of Apq12 with NPC biogenesis intermediates. In a manner that is reminiscent of yeGFP-Apq12 behaviour (Fig. 8A and B), yeGFP-Apq12-ah co- localized with Nup85-tdTomato in *td-brl1* and *td-brr6,* but not WT cells (Fig. 8D and E). The yeGFP-Apq12-ah and Nup85-tdTomato signals along the NE correlated in *td-brr6* and *td-brl1* cells but not in the WT control cells (Fig. 8F). This together suggests that Apq12 associates with NPC assembly intermediates in *td-brl1* and *td-brr6* cells and that the AαH of Apq12 is not required for this localization.

**Figure 8.**
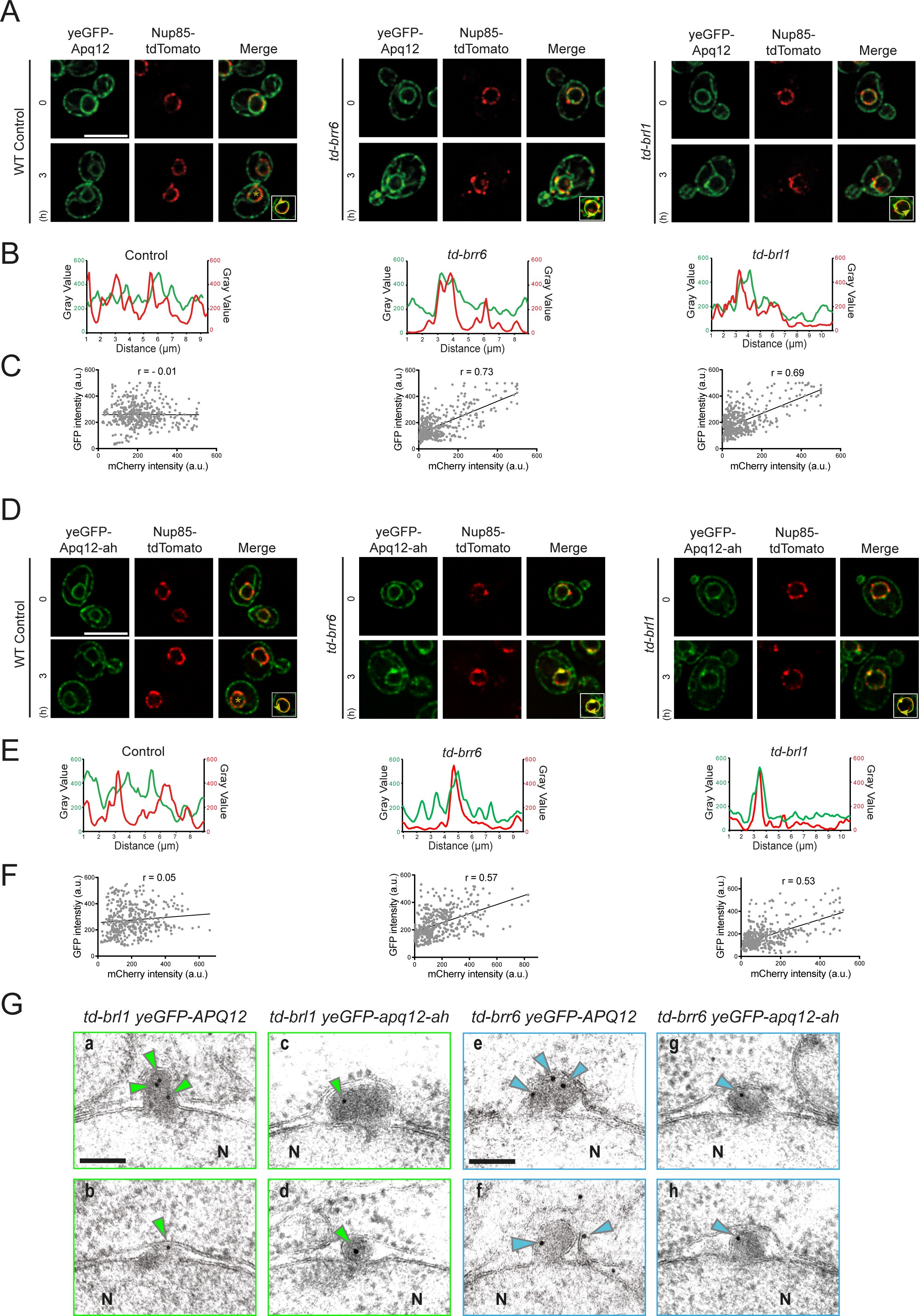
Apq12 associates with NPC biogenesis intermediates. (A) yeGFP-Apq12 localization in WT, *td-brr6* and *td-brl1* cells carrying the NPC marker *NUP85-tdTomato*. Cells were incubated for 0 h and 3 h with PGal1-*UBR1* induction at 37°C. Line scans along the NE of cells shown in the insets in (A). (C) Scatterplot with Pearson correlation coefficient (r) of GFP and mCherry fluorescence intensities (in arbitrary units, a.u.) along the nuclear rim of 5 cells. (D) yeGFP-Apq12-ah localization in WT, *td-brr6* and *td-brl1* cells carrying the NPC marker *NUP85-tdTomato*. Cells were incubated for 0 h and 3 h with PGal1-*UBR1* induction at 37°C. (E) Line scans along the NE of cells shown in the insets in (D). (F) Scatterplot with Pearson correlation coefficient (r) of GFP and mCherry fluorescence intensities (in arbitrary units, a.u.) along the nuclear rim of 5 cells. (A, D) Scale bars: 5 µm. (G) *td-brl1* and *td-brr6* cells with *yeGFP-APQ12* or *yeGFP-apq12-ah* were incubated for 3 h under restrictive conditions (galactose, 37°C) to induce degradation of Brl1 and Brr6 and the formation of herniations. Fixed and embedded cells were analyzed for localization of yeGFP- Apq12 and yeGFP-Apq12-ah by immuno-EM using GFP antibodies and 10 nm gold labelled protein A. Blue arrowheads indicate the localization of 10 nm gold particles. Scale bars: 100 nm.

In order to understand the precise localization of yeGFP-Apq12 and yeGFP-Apq12-ah in *td- brl1* and *td-brr6* cells in greater depth, immuno-EM analysis with anti-GFP antibodies was used to determine the subcellular distribution of these proteins. yeGFP-Apq12 was detected at, or inside, herniations (Fig. 8G and Fig. S7A). yeGFP-Apq12-ah was also recruited to herniations (Fig. 8E and Fig. S7A and B). To ensure that the Apq12 enrichment was not an indirect consequence of the increase in membrane surface in herniations compared to the NE, we measured gold particle number relative to the length of the sectioned membrane along NE and in herniations. This analysis detected less than 1 gold particle per µm of the NE in WT, *td-brl1* and *td-brr6* cells (Fig. S7C). In herniations, this number was increased to 5 gold particles per µm clearly indicating an enrichment of Apq12 with herniations.

### Interaction of Brl1 and Brr6 is regulated by AαH of *APQ12*

Apq12 shows physical and genetic interactions with *BRL1* and *BRR6* (Hodge et al., 2010, Lone et al., 2015, Scarcelli et al., 2007). A dysfunctional AαH could have an impact on Brl1 and Brr6 behaviour and their interaction with Apq12. Analysis of the Brl1 protein indicated an increase of Brl1 levels in *apq12Δ* and *apq12-ah* mutants (Fig. 9A and B) compared to WT. This increase was more pronounced in *apq12Δ* mutants (Fig. 9A and B). Because of the lack of Brr6 antibodies, we tested the abundance of Brr6-yeGFP in WT, *apq12Δ* and *apq12-ah* mutants. *apq12Δ* showed lethality in combination with *BRR6-yeGFP* and was therefore not tested. The Brr6-yeGFP level was increased in apq12-ah mutant compared to the WT (Fig. 9C and D). Analysis of the localization of Brl1-yeGFP and Brr6-yeGFP in *apq12-ah* mutants showed cellular distributions of both proteins that were reminiscent of that seen in WT (Fig. 9E), with an increase in signal intensity along the NE and cortical ER in *apq12-ah* mutants (Fig. 9E, Grey Value of the NE scans). This is consistent with the increase in protein levels of Brl1 and Brr6 in *apq12-ah* cells (Fig. 9A-D).

**Figure 9.**
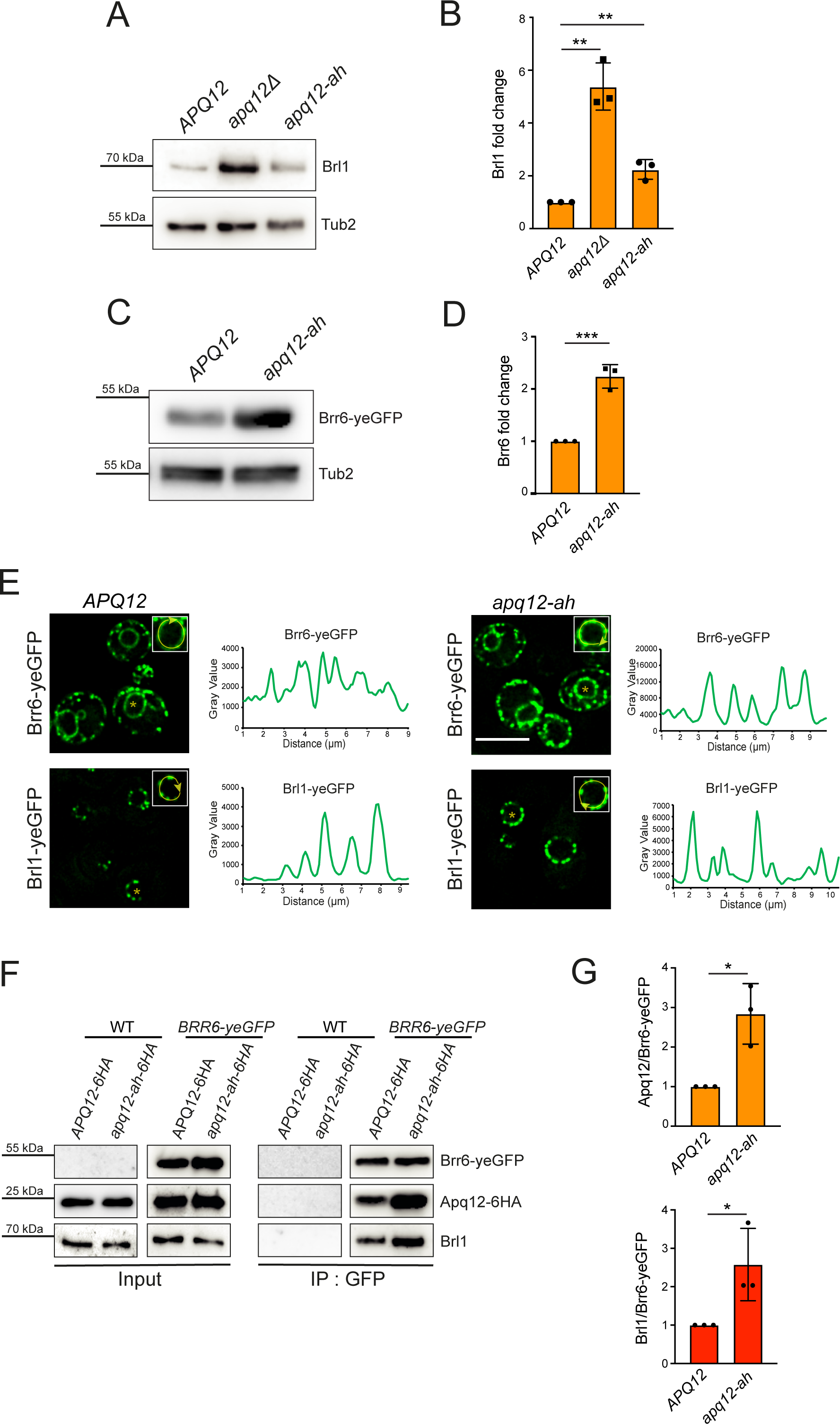
Interaction of Brl1 and Brr6 is regulated by AαH of *APQ12*. (A) Immunoblot showing endogenous levels of Brl1 in WT *APQ12*, *apq12Δ* and *apq12-ah* backgrounds, using anti-Brl1 antibody. Tub2 is loading control. (B) Quantification of Brl1 from (A) normalized to Tub2. Error bars are SD, n = 3. **p<0.01. (C) Immunoblot showing levels of Brr6-yeGFP in WT (*APQ12*) and *apq12-ah* backgrounds using anti-GFP antibody. Tub2 is loading control. (D) Quantification of Brr6 from (C) normalized to Tub2. Error bars are SD, n = 3. ***p<0.001. (E) Localization of Brr6-yeGFP and Brl1-yeGFP in WT *APQ12* (left) and *apq12-ah* cells (right) and corresponding line scans along NE of cells shown in the insets. Scale bar: 5 µm. (F) Co-IP of Apq12, Brl1 and Brr6. Brr6-yeGFP was immunoprecipitated with GFP antibodies. Apq12-6HA was detected with anti-HA and Brl1 with Brl1 antibodies. (G) Quantification of the ratio between Apq12 and Brr6-yeGFP and, Brl1 and Brr6-yeGFP in WT *APQ12* and *apq12-ah* backgrounds. Error bars are SD, n = 3; t-test; *p<0.05.

We next analyzed whether the increase in protein levels of Brl1 and Brr6 in mutants is also reflected in Brl1-Brr6 interaction in co-IP experiments in which Brr6-yeGFP was immunoprecipited with anti-GFP antibodies followed by the analysis of the immuno- precipitated proteins with Brl1 and Apq12-6HA antibodies. The Brr6-yeGFP immunoprecipitation showed complex formation of Apq12 and Apq12-ah with Brl1 and Brr6 (Fig. 9F). Quantification of the immunoprecipitated proteins from three independent experiments showed that although Brr6-yeGFP precipitation efficiency was very similar, more Apq12 and Brl1 were co-immunoprecipitated from *apq12-ah* cells than from *APQ12* cells (Fig. 9F and G). Thus, a defect in AαH of Apq12 enhances the interaction between Brr6, Brl1 and Apq12.

## Discussion

Apq12 is important for NPC function and cooperates with the essential NPC biogenesis factors Brl1 and Brr6 (Hodge et al., 2010, Lone et al., 2015, Scarcelli et al., 2007). These data indicate that Apq12 is at the heart of machinery that ensures NPC assembly, raising the important question about its molecular function. Because Apq12 does not have any sequence similarities to proteins with enzymatic activity, we have to assume that it functions either as a protein scaffold or impacts the shape of the NE as reported for reticulons that bend the ER by two cooperating pairs of trans-membrane domains with an adjacent AαH (Hu, Shibata et al., 2008, Wang, Clark et al., 2021). Following the rational that a combination of transmembrane domains and AαH could be the basis for the functional of Apq12, we analyzed Apq12 by the AmphipaSeeK program that predicated the presence of a short AαH connecting the two transmembrane domains. Consistent with this predication, the synthetic AαH peptide binds to liposomes dependent on its amphipathic nature and the lipid composition. In addition, topological analysis showed that the AαH resides in the intermembrane space of the NE, while N- and C-terminal regions of Apq12 localize in the nucleus or cytoplasm depending on the INM or ONM localization of Apq12.

In terms of cold sensitivity, NE and NPC biogenesis defects the *apq12-ah* mutant behaves similar to complete loss of *APQ12* function (this study, (Hodge et al., 2010, Lone et al., 2015, Scarcelli et al., 2007). However, phenotypic differences between *apq12-ah* and *apq12Δ* are also detectable (Fig. 3). In addition, Apq12-ah associates at least as efficiently with Brl1 and Brr6 as Apq12, has the same topological arrangement as Apq12 and associates with NPC assembly intermediates. This together indicates that *apq12-ah* is not a loss of function allele. The AαH is required for the overall function of Apq12. However, N- and C-termini of Apq12 fulfill functions without the involvement of AαH. This probably explains why prolonged overexpression of *apq12-ah* still shows a mild membrane deforming phenotype (Fig. 5).

*apq12-ah* and *apq12Δ* mutants impact cellular lipid composition, independent of the tested growth temperature, in a way that reduces membrane fluidity. This accounts for the cold sensitive growth defect and the NE breakdown phenotype at reduced growth temperatures. In addition, we observed the striking accumulation of PA either through synthesis or re- localization at the NE upon overexpression of Apq12, dependent on the functionality of the AαH. Before, it was shown that PA accumulates at sites of NPC mis-assembly (Thaller, Tong et al., 2021) raising the question as to whether PA accumulation at the NE induced by PGal1- *APQ12* overexpression is a consequence of defective NPCs. The observation that PGal1- *APQ12* overexpression merely displaces NPCs into areas that lack ONM expansions without the accumulation of defective NPC assemblies (Figs. 4C and S3A), indicates that overproduced Apq12 has the ability to induce PA accumulation at the NE even when NPCs are intact.

Before we have shown that INM membrane deformations in *td-brl1* and *td-brr6* cells reflect emerging NPCs that do not fully assemble because of a defect in one of the steps leading to the generation of functional NPCs (Zhang et al., 2018). Using cells carrying *td-brl1* and *td- brr6* we observed strong accumulation of Apq12 with NPC biogenesis intermediates and this localization does not require a functional AαH. In contrast, Apq12 only co-localizes with a small number of NPCs in WT cells. Thus, Apq12 joins Brl1 and Brr6 (Zhang et al., 2018) in a small subset of proteins required for NPC assembly that transiently interact with assembling NPCs but then dissociate from NPCs as soon as they are fully assembled.

As Apq12, Brl1 and Brr6, the Lap2-emerin-MAN1 (LEM) family proteins Heh1 and Heh2 are also nonstructural components of NPCs. However, in contrast to Apq12, Brl1 and Brr6 that function directly in NPC biogenesis, Heh1 and Heh2 contribute to the surveillance of NPC biogenesis (Webster, Colombi et al., 2014). Upon NE damage caused by a NPC biogenesis defect, Chm7, a component of an ESCRT-III like complex, enters the nucleus where it is activated by nuclear Heh1 to promote the sealing of the disrupted NE by the ESCRT machinery (Thaller et al., 2019, Webster et al., 2014). In contrast to Heh1, Heh2 probably functions as a senor for the assembly state of NPCs (Borah, Thaller et al., 2021). Thus, Apq12, Brl1 and Brr6 and Heh1 and Heh2 fulfill quite distinct functions at NPCs.

Based on our findings and the observation that the role of the Apq12-Brl1/Brr6 module overlaps with that of the nucleoporin Nup116 in scaffolding NPC biogenesis (Onischenko et al., 2017, Zhang et al., 2018), we suggest the following stages for the stepwise assembly of NPC assembly. The FG nucleoporin Nup116 was shown to scaffold NPC assembly together with Nup188 on the nuclear side of the INM (Onischenko, Noor et al., 2020, Onischenko et al., 2017). As Nup116, Apq12, Brl1 and Brr6 also associate with NPC intermediates (van Leeuwen, Pons et al., 2020, Zhang et al., 2018 and this study). The interaction of Brl1 with the integral membrane protein Ndc1 (Winey, Hoyt et al., 1993) and Nup188 could be the trigger for the recruitment of this protein to assembling NPCs (Zhang et al., 2018). How Apq12 is recruited to NPC intermediates is not understood. The interacting Brl1 and Brr6 are not essential for this localization since Apq12 associates with herniations upon induced depletion of Brl1 and Brr6 (Fig. 8).

Apq12 probably executes multiple functions, some of which are at least partly mediated by its AαH at NPC biogenesis sites. First, Apq12 inserts through its membrane active AαH into the NE from within the intermembrane space (Fig. 1). This insertion has the potential to generate membrane curvature that may stabilize membrane deformations that arise during NPC biogenesis. AαH peptide did not deform GUVs under the experimental conditions we applied (Fig. 1). However, the yeast reticulon Yop1 only deforms liposomes in the context of transmembrane domains and AαH (Wang et al., 2021). Thus, in future experiments it will be important to measure the membrane-deforming ability of the TM-AαH-TM core of Apq12. Second, the enrichment of Apq12 at NPC biogenesis sites may promote local accumulation of PA as indicated by the *APQ12* overexpression phenotype. Because of its conical shape, PA could stabilize bent membrane regions at INM bends during NPC biogenesis (Zhukovsky, Filograna et al., 2019). In addition, PA has the ability to interact with a range of proteins via short stretches of positively charged amino acid residues (Jang, Lee et al., 2012, Tanguy, Kassas et al., 2018). Presently, no Nup or NPC biogenesis factor with PA binding activity has been described. However, it was only until recently that the PA-binding activity of the ESCRTIII protein Chm7 was reported (Thaller et al., 2021), raising the possibility that proteins involved in NPC assembly may carry hidden PA binding sites. Third, considering that Apq12 has an impact on Brl1 and Brr6 levels and the interaction of Apq12, Brl1 and Brr6 (Fig. 9), it may coordinate localization of the interacting Brl1 and Brr6 at NPCs biogenesis sites, a function that would be most important at elevated temperatures when Apq12 most strongly plays a role in NPC assembly. *APQ12*, *BRL1* and *NUP116* show genetic interactions and overexpression of *BRL1* is able to suppress the NPC biogenesis defect of *nup116Δ* and *nup116ΔGLFG PMET3-NUP188* mutants by promoting the fusion of the INM and ONM (Onischenko et al., 2017, Scarcelli et al., 2007, Zhang et al., 2018). We therefore suggest that Brl1, in part scaffolded by Apq12, directly or indirectly, facilitates the fusion between the INM with the ONM during NPC biogenesis.

The Apq12-Brl1/Brr6 module is only conserved in organisms with closed mitosis (Tamm et al., 2011). However, considering the simple domain architecture of Apq12 with two TM domains and a short AαH, proteins with similar organisation may substitute for the function of Apq12 in higher eukaryotes without having detectable amino acid homology to Apq12, rather a structural similarity with two closely opposed TM domains. Brl1 and Brr6 have an extended intermembrane space domain that is stabilized by two functionally important disulphide bridges (Tamm et al., 2011, Zhang et al., 2018). This stabilization principal could be substituted for alternative structural features in functional equivalent proteins in higher eukaryotes.

## Materials and Methods

### Yeast strains and plasmids

Yeast strains and plasmids used in this study are listed in Table S1. All yeast strains are derived from ESM356-1 (*MATa ura3-52 trp1Δ63 his3Δ200 leu2Δ1*). Gene deletion and epitope tagging of endogenous genes was performed using a PCR-based integration approach (Janke, Magiera et al., 2004, Knop, Siegers et al., 1999). Yeast strains were grown in SC (synthetic complete) medium, SC-selection medium (Guthrie & Fink, 1991), YPD (yeast extract, peptone and glucose) or YPRaf (yeast extract, peptone and raffinose) with or without 0.1 mM CuSO4 at 23°C, 30°C or 37°C. Galactose was added to a final concentration of 2% to induce expression of genes under control of the PGal1 promoter. Alkaline lysis and TCA precipitations were used to prepare yeast extracts in order to analyse protein levels by immunoblotting (Janke et al., 2004). To test for growth defects, yeast cells were grown over night in the indicated selection medium, afterwards the cell density was adjusted to OD600 = 1. The cell suspension was then spotted in 10-fold serial dilutions onto selection plates that were incubated as indicated.

### EM and immuno-EM analysis of yeast cells

Cells were high pressure frozen, freeze-substituted, sectioned, labelled and stained for electron microscopy as described (Giddings, O’Toole et al., 2001). Briefly, cells were collected onto a 0.45 μm polycarbonate filter (Millipore) using vacuum filtration and then high pressure frozen with a HPM010 (Abra-Fluid, Switzerland). Cells were freeze-substituted using the EM-AFS2 device (Leica Microsystems, Vienna, Austria) (freeze substitution solution: 0.1% glutaraldehyde, 0.2% uranyl acetate, 1% water – dissolved in anhydrous acetone) and stepwise infiltrated with Lowicryl HM20 (Polysciences, Inc., Warrington, PA), started at -90°C. For polymerization the samples were exposed to UV light for 48 h at -45°C and gradually warmed up to 20°C. Embedded cells were serially sectioned using a Reichert Ultracut S Microtome (Leica Instruments, Vienna, Austria) to a thickness of 80 nm. Post-staining with 3% uranyl acetate and lead citrate was performed. Sections were imaged at a Jeol JE-1400 (Jeol Ltd., Tokyo, Japan) operating at 80 kV equipped with a 4k x 4k digital camera (F416, TVIPS, Gauting, Germany). Micrographs were adjusted in brightness and contrast using ImageJ. For immuno-labelling the primary antibody was used against GFP. The samples were prepared similarly with the exception that the glutaraldehyde was omitted from the freeze substitution solution. The sections were treated with blocking buffer (1.5% BSA, 0.1% fish skin gelatin in PBS), then incubated with the primary antibody, followed by treatment with protein A-gold conjugates (10 nm, Utrecht University, Utrecht, The Netherlands).

### Fluorescence light microscopy and image analysis

A DeltaVision RT system (Applied precision, Olympus IX71 based) equipped with the Photometrics CoolSnap HQ camera (Roper Scientific), a 100x/1.4 NA UPlanSAPO objective (Olympus), a mercury arc light source and the softWoRx software (Applied Precision) was used for cell imaging. Imaging was done at 16°C, 23°C, 30°C or 37°C using the GFP, and the mCherry channels with different exposure times according to the fluorescence intensity of each protein. For time-lapse experiments, cells were immobilized on Concanavalin A (Sigma-Aldrich)–coated 35-mm glass bottomed dishes (P35G-1.5-14C; MatTek Corporation) and kept in their respective media. Images were deconvolved with the softWoRx software (Applied Precision) and processed with ImageJ (National Institutes of Health, Bethesda MD). For Fig. 5 C and E, Quantification of maximum intensity of individual cells was done by utilising CellProfiler 3.1.9 software (Broad Institute, Cambridge, MA) (Carpenter, Jones et al., 2006).

### Lipid analysis by mass spectrometry

10 OD of cells were harvested and homogenized using a FastPrep machine (MP Biomedicals) in 155 mM ammonium bicarbonate buffer at pH 7.5. Homogenized cells were subjected to acidic Bligh&Dyer lipid extractions in the presence of internal lipid standards added from a master mix containing PC (phosphatidylcholine, 13:0/13:0, 14:0/14:0, 20:0/20:0; 21:0/21:0, Avanti Polar Lipids), PI (phosphatidylinositol, 17:0/20:4, Avanti Polar Lipids), PE and PS (phosphatidylethanolamine and phosphatidylserine, 14:1/14:1, 20:1/20:1, 22:1/22:1, semi-synthesized as described (Ozbalci, Boyaci et al., 2013), DAG (diacylglycerol, 17:0/17:0, Larodan), TAG (triacylglycerol, D5-TAG-Mix, LM-6000/D5-TAG 17:0,17:1,17:1, Avanti Polar Lipids), PA (phosphatidic acid, 17:0/20:4, Avanti Polar Lipids), PG (phosphatidylglycerol, 14:1/14:1, 20:1/20:1, 22:1/22:1, semi-synthesized (Ozbalci et al., 2013), and Cer (ceramide, Avanti Polar Lipids). Lipids recovered in the organic extraction phase were evaporated by a gentle stream of nitrogen. Prior to measurements, lipid extracts were dissolved in 10 mM ammonium acetate in methanol and transferred to 96-well plates (Eppendorf twintec plate 96). Mass spectrometric measurements were performed in positive ion mode on an AB SCIEX QTRAP 6500+ mass spectrometer, equipped with chip-based (HD-D ESI Chip, Advion Biosciences) nano-electrospray infusion and ionization (Triversa Nanomate, Advion Biosciences) as described (Ozbalci et al., 2013). The following precursor ion scanning (PREC) and neutral loss scanning (NL) modes were used for the measurement of the various lipid classes: +PREC 184 (PC), +PREC282 (t-Cer), +NL141 (PE), +NL185 (PS), +NL277 (PI), + NL189 (PG), +NL115 (PA), +PREC 77 (ergosterol), + PREC379 (ergosteryl ester). Ergosterol was quantified following derivatization to ergosterol acetate in the presence of the internal standard (22E)-Stigmasta-5,7,22-trien-3-beta-ol (Aldrich, R202967) using 100 µl acetic anhydride/chloroform (1:12 v/v) (Ejsing, Sampaio et al., 2009). Data evaluation was done using LipidView (ABSciex) and an in house-developed software (ShinyLipids).

### Liposome preparation

All lipids were received from Avanti Polar lipids with the exception of Atto647N, which was obtained from Atto-Tec. The lipid composition of the PM mix consisted of: 34.8 mol% 1-palmitoyl-2-oleoyl-*sn*-glycero-3-phosphocholine (POPC), 15 mol% 1,2-dioleoyl-*sn*-glycero- 3-phosphoserine (DOPS), 20 mol% 1-hexadecanoyl-2-octadecenoyl-*sn*-glycero-3-phospho- ethanolamine (POPE), 25 mol% cholesterol (from ovine wool), 5 mol% liver L-α- phosphatidylinositol (PI, from liver) and 0.2 mol% Atto647N-DPPE. The composition of the used nuclear envelope (NE) lipid mix was: 19.8 mol% POPC, 3 mol% DOPS, 42 mol % Cholesterol, 7 mol% POPE, 23 mol % PI, 5 mol% PI(4,5)P2, and 0.2 mol% Atto647 (Zhendre, Grelard et al., 2011). SUVs (small uni-lamellar vesicles) were formed as described previous (Weber, Zemelman et al., 1998) by dissolving the lipid mixtures in Octyl-β-D glucopyranoside (OG) containing buffer, OG dilution below the critical micellar concentration, flow dialysis and SUV isolation using Nycodenz-gradient centrifugation. Subsequently, the concentrated liposomes were extruded 23-times with a 100 nm filter and stored at 4°C.

For preparation of GUVs (giant uni-lamellar vesicles) the PM mix was used. SUVs were prepared to produce GUVs as described previously (Malsam, Parisotto et al., 2012) with the following modifications: (I) the GUVs were desalted two times employing a PD10 column (GE Healthcare) instead of using a Sephadex-G50 gel filtration column in the second desalting step. (II) platinum-coated glass slides (GeSiM) were applied instead of ITO-coated glass slides (GeSiM) (Meijer, Dorr et al., 2018).

The lipid concentration was measured by the fluorescence of Atto647N with an excitation of 647 nm and an emission of 670 nm using a Fluoroskan Ascent FL plate reader (Thermo Scientific) in a 96-well plate (integration time: 1000 ms). Therefore, the liposomes were disrupted by 0.5% Dodecyl-maltoside (DDM). The recovery of the total lipid was compared to the lipid after preparation.

### Dynamic light scattering (DLS) of the SUVs

DLS was applied to monitor the size of the liposomes. After adding 2 µM lipid (5 µl final volume) to the quartz cuvette, the particle size was determined in a DynaPor NanoStar (Wyatt technologies) instrument at room temperature. The buffer composition for size determination was set to PBS and 20 acquisitions with 5 s time intervals were measured.

### Binding studies of SUVs

The binding of the AαH and AαH-ah peptides (from PSL Peptide Specialty Laboratories GmbH, Heidelberg; amino acid sequence: KLLMNFITLVKRFL, KLDMNRNTLVKRNL) coupled with Atto488 (freshly solved in DMSO and diluted in 25 mM HEPES pH 7.4, 135 mM KCl, 1 mM DTT (fusion buffer)) with SUVs was measured by a Nycodenz gradient centrifugation. 0.45 mM lipid was incubated with 18 µM AαH or AαH-ah peptide or 18 µM Atto488 in 25 mM HEPES pH7.4, 135 mM KCl, 2.1% DMSO and 1 mM DTT for 30 minutes at room temperature, whereas 10% of the sample was used as the reference input. The peptide without liposomes functioned as a floatation control. Nycodenz (solved in fusion buffer) was added to the sample producing a 40% Nycodenz layer in the bottom of the centrifuged tube. The sample was overlaid with decreasing concentrations of Nycodenz (30%, 20%, 10% and 0%) and centrifuged in a SW55 rotor (Beckman Coulter) 2h 30 min at 55000 rpm and 4°C. The floated SUVs were located in the layer between 10% – 20% Nycodenz. The recovery of the SUVs was measured by the fluorescence of Atto647N in the DDM lysed sample and compared to the input. The binding of the peptide or Atto488 was analyzed by the fluorescence signal of Atto488 (excitation: 460 nm, emission: 538 nm, integration time: 1000 ms) in this layer and compared to the input probe. The binding of the peptide was corrected to the amount of the floated SUVs.

### Binding studies of GUVs

For the peptide interaction to GUVs, 3.3 µM GUVs were incubated with different amounts of peptide in a ratio of 1:1 to 10:1 in fusion buffer at room temperature for 15 minutes. As a control analysing the change of the GUV morphology 0.04% DMSO was added to the GUVs without peptide. The morphology of the GUVs was visible in the fluorescence of Atto647 and the interaction of the peptide by the fluorescent signal of Atto488. It was measured in a chambered coverslip (Ibidi, catalogue No. 80826) using a DeltaVision microscope.

### MST

For determination of the KD values via MST 2 µM Atto488 coupled to AαH and AαH-ah peptide or Atto488 as a control was incubated 15 minutes at room temperature with increasing liposome concentrations in fusion buffer with 0.01% Triton X-100. All samples were filled in Premium capillaries and the thermophoresis was measured in a Monolith NT.115 (NanoTemper) instrument with a LED power of 40% (green filter) and an MST power of 17%. The MST data were evaluated using a single exponential function in consideration of the initial state calculated by linear regression. The data were normalized to the ratio of bound/unbound peptide and the KD value was determined according to the Hill equation:

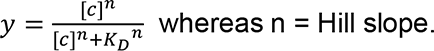

### Immunoprecipitation

Cells (25 OD) were harvested and resuspended in lysis buffer (20 mM Tris-Cl, pH 8.0, 150 mM NaCl, 5 mM MgCl2, and 10% glycerol) supplemented with 10 mM NaF, 60 mM β-glycerophosphate, 1 tablet/50 ml Roche protease inhibitor cocktail complete (EDTA free), and 1 mM PMSF. Glass beads (BioSpec Products) were added, and cells were lysed in a FastPrep machine (MP Biomedicals). Cell lysate was supplemented with 0.5% Triton X-100 and incubated on ice for 10 min. The soluble proteins were separated from the cell debris by centrifugation and incubated with GFP-Trap agarose beads (Chromotek) at 4°C for 2 h. Beads were washed three times with lysis buffer supplemented with 0.1% Triton X-100 and twice with wash buffer (20 mM Tris-Cl, pH 8.0, 150 mM NaCl, and 5 mM MgCl2). Bound proteins were eluted in 50 µl of 2× SDS-PAGE sample buffer, heated to 95°C for 5 min, separated by SDS-PAGE, and transferred to PVDF membrane (Millipore) for Western blotting.

### Statistical analysis

For the statistical analyses, PRISM v.7 software (GraphPad) was used. Comparisons of samples were performed using unpaired *t* test with two-tailed p-value. Data distribution was assumed to be normal, but this was not formally tested.

### Antibodies

Antibodies and their conditions of use are as follows: mouse anti-His (Western blot, 1:1000; 34660; Qiagen), rabbit anti-Brl1 (Western blot, 1:1000; made in-house), rabbit anti-Tub2 (Western blot, 1:1000; made in-house), and rabbit anti-GFP (Western blot, 1:1000; Proteintech), rabbit anti-HA (Western blot, 1:500; Proteintech), rabbit anti-GFP (immuno-EM, 1:5; gift from M. Seedorf, Zentrum für Molekulare Biologie, Heidelberg, Germany).

## Acknowledgements

We thank the electron microscopy facility of Heidelberg University for their support. This work is supported by a grant of the Deutschen Forschungsgemeinschaft (DFG) to E. Schiebel (DFG Schi 295/5-3) and to B. Brügger (Project Number 112927078 - TRR 83). We thank A. Köhler for the gift of the PA-mCherry sensor plasmids and Iain Hagan for the Brl1 antibody.

## Online supplementary material

Fig. S1 shows the quantification of localization data from immuno-EM. It also includes data showing the topology of Apq12 and apq12-ah by biotinylation of the HBH tag. Fig. S2 shows the lipid analysis of WT, *apq12**Δ*** and *apq12-ah* cells. Fig. S3 shows time courses and time lapses of Apq12 overexpression under the galactose promoter in *yeGFP-BRL1 NUP85- tdTomato* and *yeGFP-BRR6 NUP85-tdTomato* cells. Fig. S4 shows effects of *APQ12* overexpression on the INM protein Asi3 and ER luminal marker HDEL. Fig. S5 shows the localization of overexpressed Apq12-yeGFP and apq12-ah-yeGFP with cellular sites of membrane over proliferation. Fig. S6 shows the lipid analysis of cells with overexpression of Apq12 and Apq12-ah. Fig. S7 shows immuno-EM images of yeGFP-Apq12 and apq12-ah- yeGFP localizing with herniations.

## Author Contributions

Wanlu Zhang

Zentrum für Molekulare Biologie der Universität Heidelberg, DKFZ-ZMBH Allianz, Heidelberg, Germany.

Cell Biology and Biophysics Unit, European Molecular Biology Laboratory (EMBL), Heidelberg, Germany.

**Contribution:** Conceptualization, Investigation, Formal analysis, Visualization, Review and editing

**Competing Interest:** No competing interests declared

Azqa Khan

Zentrum für Molekulare Biologie der Universität Heidelberg, DKFZ-ZMBH Allianz, Heidelberg, Germany.

**Contribution:** Investigation, Formal analysis, Data curation and visualization, Review and editing

**Competing Interest:** No competing interests declared

Jlenia Vitale

Zentrum für Molekulare Biologie der Universität Heidelberg, DKFZ-ZMBH Allianz, Heidelberg, Germany.

**Contribution:** Investigation, Formal analysis and visualization

**Competing Interest:** No competing interests declared

Annett Neuner

Zentrum für Molekulare Biologie der Universität Heidelberg, DKFZ-ZMBH Allianz, Heidelberg, Germany.

**Contribution:** EM and immune-EM and analysis

**Competing Interest:** No competing interests declared

Kerstin Rink

Biochemie-Zentrum der Universität Heidelberg, Heidelberg, Germany **Contribution:** Design and analysis of lipid binding experiments **Competing Interest:** No competing interests declared

Christian Lüchtenborg

Biochemie-Zentrum der Universität Heidelberg, Heidelberg, Germany

**Contribution:** Lipid mass spectrometry analysis

**Competing Interest:** No competing interests declared

Britta Brügger

Biochemie-Zentrum der Universität Heidelberg, Heidelberg, Germany

**Contribution:** Lipid mass spectrometry analysis

**Competing Interest:** No competing interests declared

Thomas H. Söllner

Biochemie-Zentrum der Universität Heidelberg, Heidelberg, Germany **Contribution:** Design and analysis of lipid binding experiments **Competing Interest:** No competing interests declared

Elmar Schiebel

Zentrum für Molekulare Biologie der Universität Heidelberg, DKFZ-ZMBH Allianz, Heidelberg, Germany.

**Contribution:** Conceptualization, Supervision, Funding acquisition, Writing, Review and editing

**Competing Interest:** No competing interests declared

---

**Figure S1.**
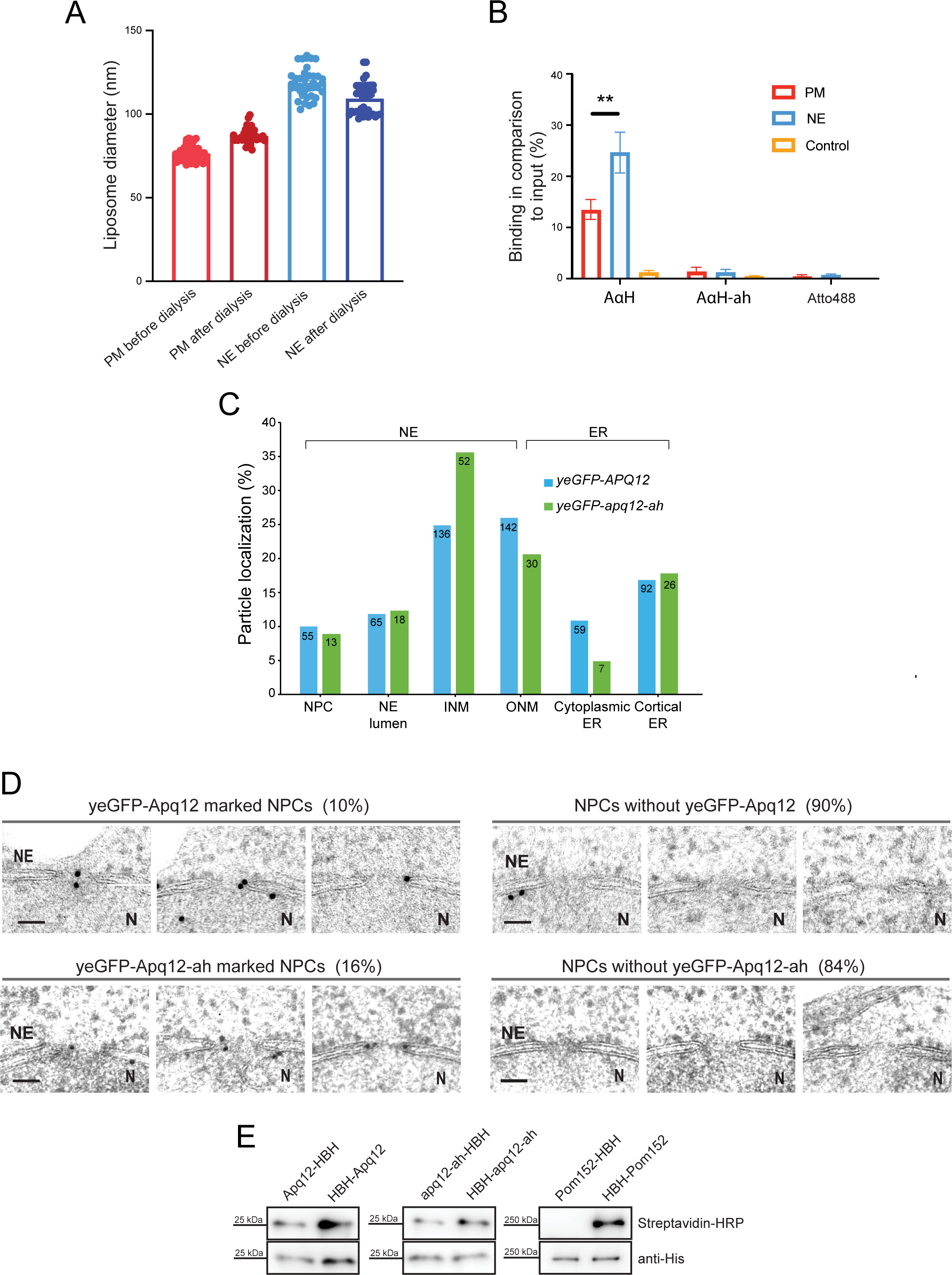
Localization and topology of Apq12 and Apq12-ah. (A) The diameter of liposomes of PM or NE lipids was measured by DLS with 20 acquisitions at room temperature before and after dialysis with the MST buffer. (B) Binding of AαH and AαH-ah peptides to liposomes measured by float-up. The data were compared to the input. Values are given as means ± S.E.M (n = 3; t-test: **p<0.01). (C) Quantification of data from Fig. 2C. Number of analyzed gold particles for each category is given in the figure. (D) Immuno-EM images of cells with and without yeGFP-Apq12 and yeGFP-Apq12-ah at NPCs. For yeGFP-Apq12 and yeGFP-Apq12-ah 12 and 33 NPCs were analyzed. Scale bars: 50 nm. (E) Analysis of the membrane topology of Apq12. The indicated yeast cells were analyzed by immunoblotting for the biotinylation of the HBH tag for cytoplasmic or nuclear localization. N and C terminally tagged Pom152 is used as a control.

**Figure S2.**
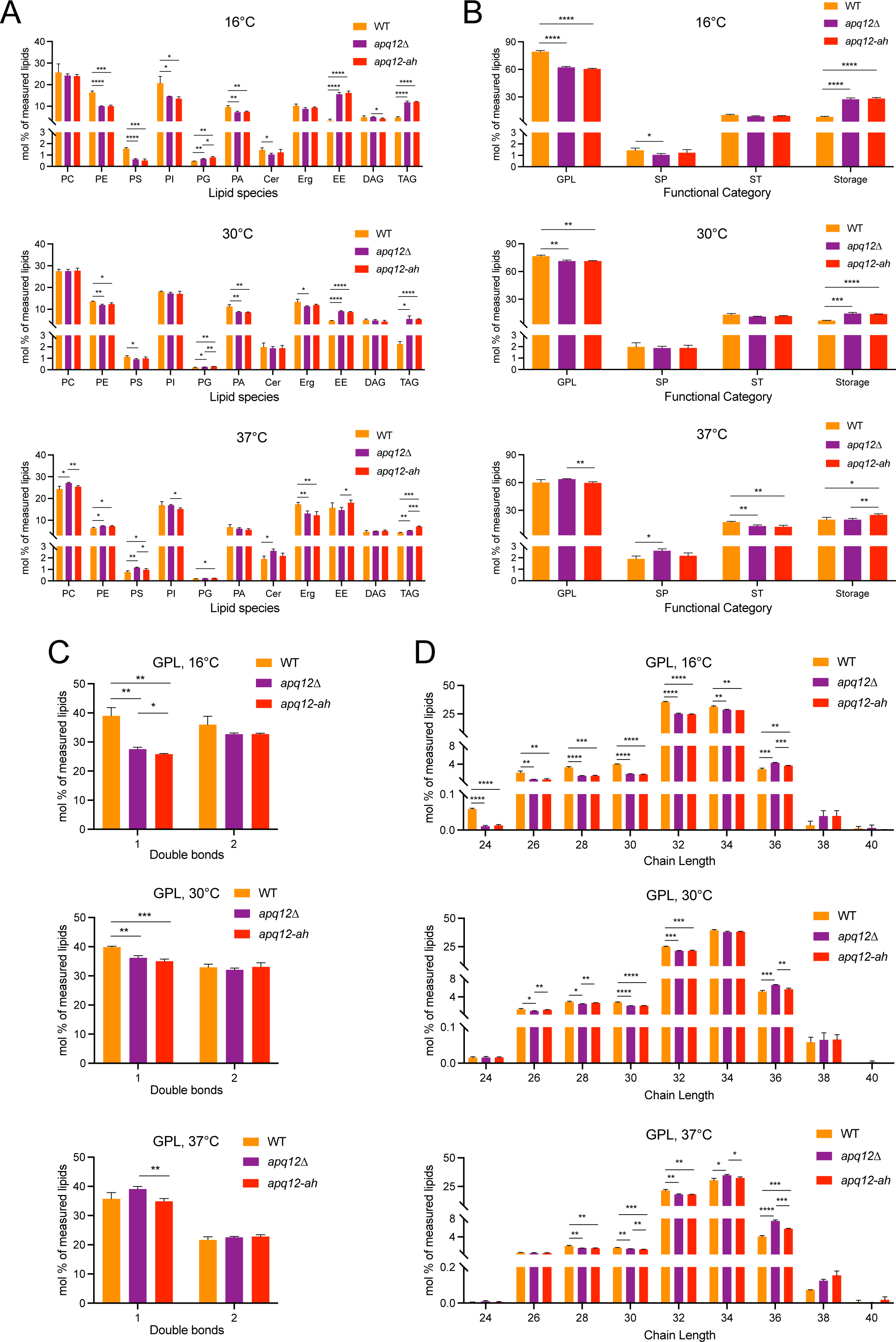
Lipid analysis of WT and *apq12Δ* cells. Lipids were extracted from *APQ12* WT, *apq12Δ* and *apq12-ah* cells incubated at the indicated temperatures and subsequently analyzed by nano-ESI-MS/MS. Samples were taken after 2 h at 37°C, 4 h at 30°C and 18 h at 16°C. (A) Plots of lipid species in mol% of measured lipids. (B) Plots of functional categories of lipids; glycerophospholipids and glycerolipids (GPL), sphingolipids (SP), sterols (ST), and storage lipids. (C) Plots showing the number of double bonds in GPL. (D) Plots showing the chain lengths of the GPL. n = 3. Values are given as means + SD. t-test *p<0.05, **p<0.01, ***p<0.001, ****p<0.0001.

**Figure S3.**
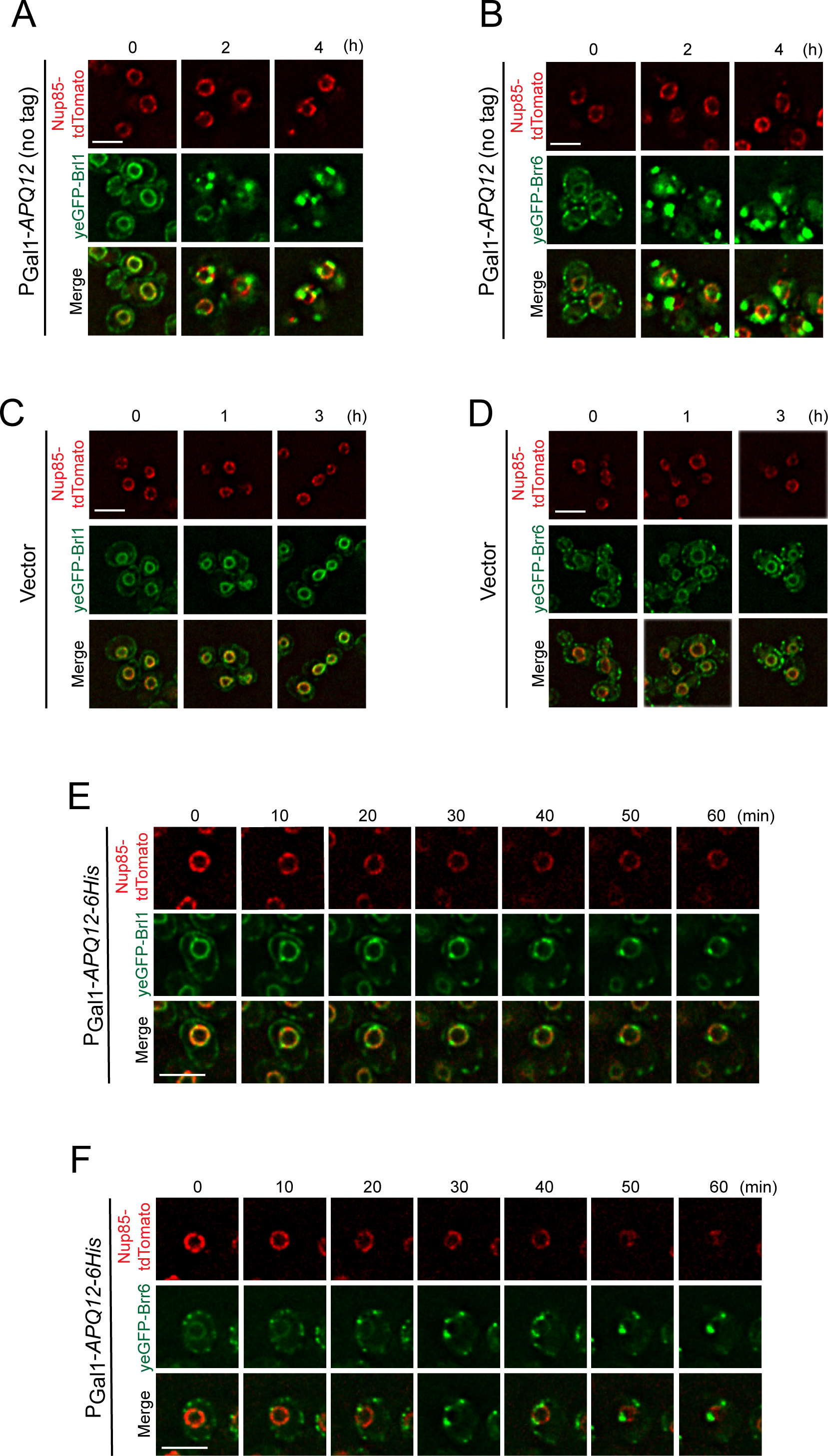
Overexpression of *APQ12-6His* has a similar impact as that of *APQ12*. (A, B) Induction of PGal1-*APQ12* caused the same phenotypes in *yeGFP-BRL1 NUP85- tdTomato* (A) and *yeGFP-BRR6 NUP85-tdTomato* (B) cells as seen for PGal1-*APQ12-6His*. (C, D) Cells with the vector control p425-PGal1 show no impact on yeGFP-Brl1, yeGFP-Brr6 and Nup85-tdTomato upon addition of galactose (t=0). (E) Live-cell imaging of yeGFP-Brl1 and Nup85-tdTomato with induction of PGal1-*APQ12* by addition of galactose (t=0). (F) Live-cell imaging of yeGFP-Brr6 and Nup85-tdTomato with induction of PGal1-*APQ12* by addition of galactose (t=0). (A-F) Scale bars: 5 µm.

**Figure S4.**
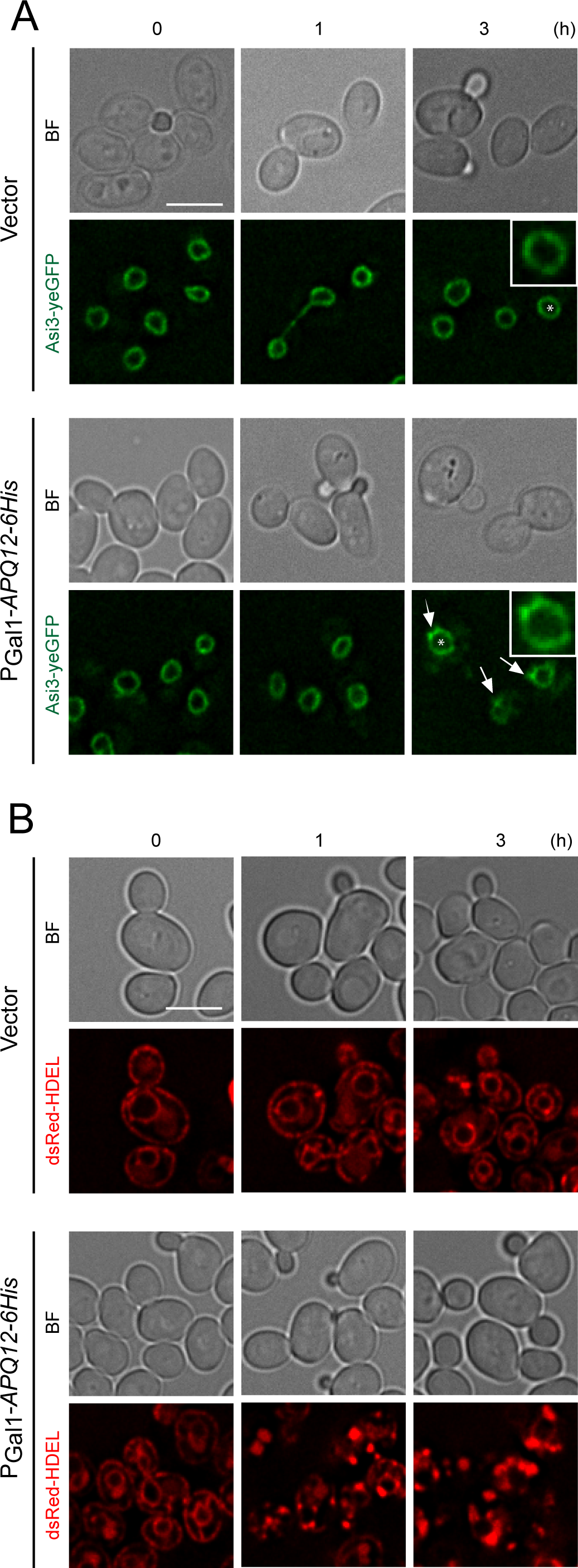
Overexpression of *APQ12-6His* weakly affects localization of the INM protein Asi3 but strongly affects the ER luminal marker HDEL. (A, B) Impact of PGal1-*APQ12* overexpression on the INM protein Asi3-yeGFP (A) and the ER marker dsRed-HDEL (B). Induction of PGal1-*APQ12* was triggered by the addition of galactose (t=0). The arrows in (A) indicate small enrichments of Asi3-yeGFP at the NE after 3 h of PGal1-*APQ12-6His* induction. Insets in (A) are two-fold enlargements of a cell in the overview marked by an asterisk. (A, B) BF: brightfield. Scale bars: (A, B) 5 µm.

**Figure S5.**
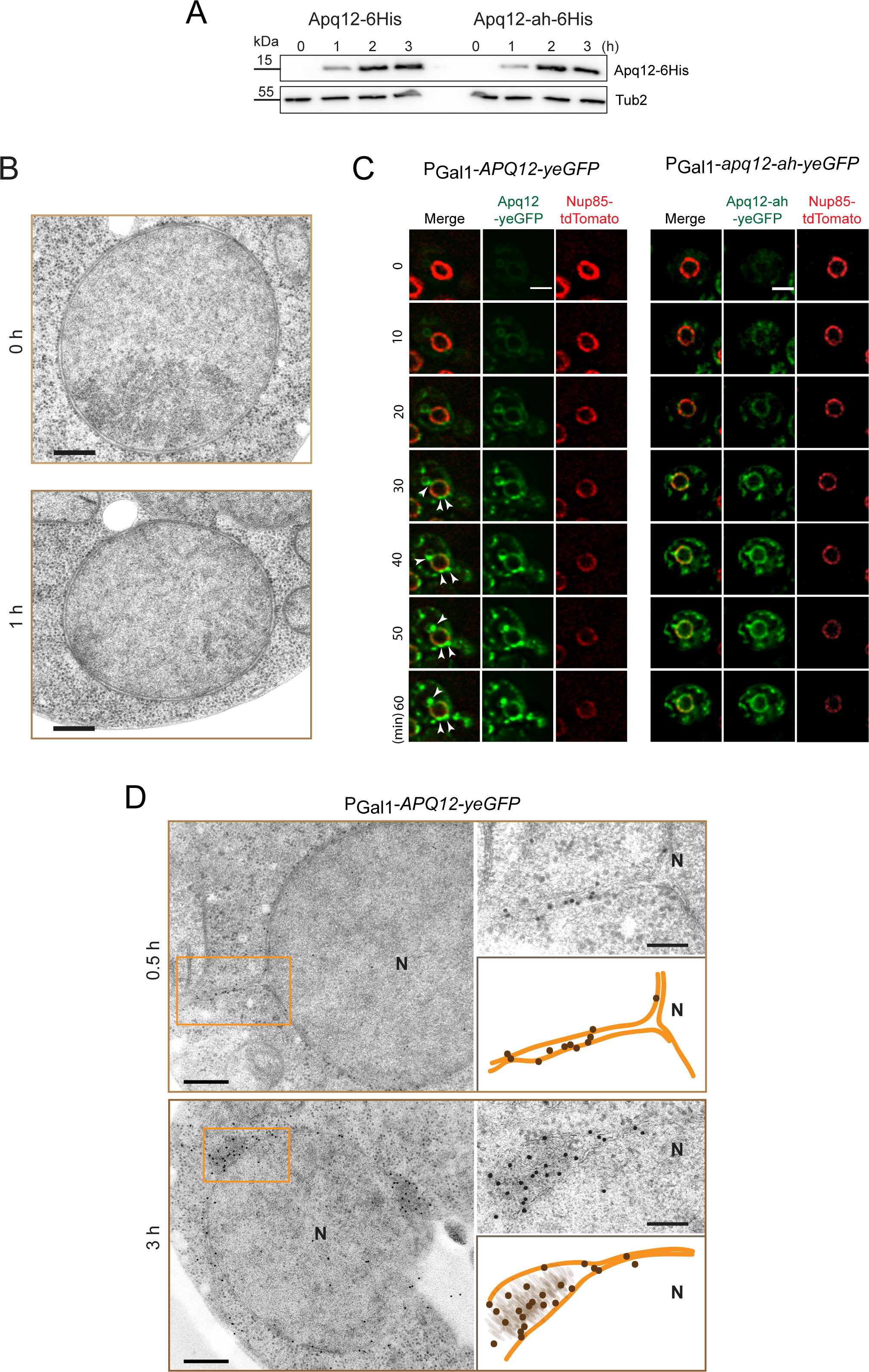
Apq12-yeGFP localizes with cellular sites of membrane over proliferation. (A) Immunoblot showing induction of PGal1-*APQ12-6His* and PGal1*-apq12-ah-6His* after addition of galactose. His antibodies detected Apq12-6His and Apq12-ah-6His. Tub2 is loading control. (B) EM analysis of PGal1-*APQ12-6His* cells at t=0 and vector control pGal1 cells at t=1 of galactose addition. Scale bars: 250 nm. (C) Live-cell imaging of Apq12-yeGFP (left) and Apq12-ah-yeGFP (right) in combination with Nup85-tdTomato upon galactose (t=0) induced overexpression of PGal1-*APQ12-yeGFP* and PGal1-*apq12-ah-yeGFP*, respectively. White arrowheads indicate NE extensions. Scale bars: 3 µm. (D) Immuno-EM localization of Apq12-yeGFP in cells expressing PGal1-*APQ12-yeGFP* for 0.5 and 3 h induction by addition of galactose. The 10 nm gold particles indicate localization of Apq12-yeGFP. The ochre rectangles on the left illustrate the magnifications that are shown on the upper right. The cartoons on the lower right are schematic views on the ER/ONM of the EM pictures in the top right. Gold particles indicating the localization of Apq12-yeGFP are shown in the cartoons. Abbreviation: N, nucleus. Scale bars: 250 nm and enragements 50 nm.

**Figure S6.**
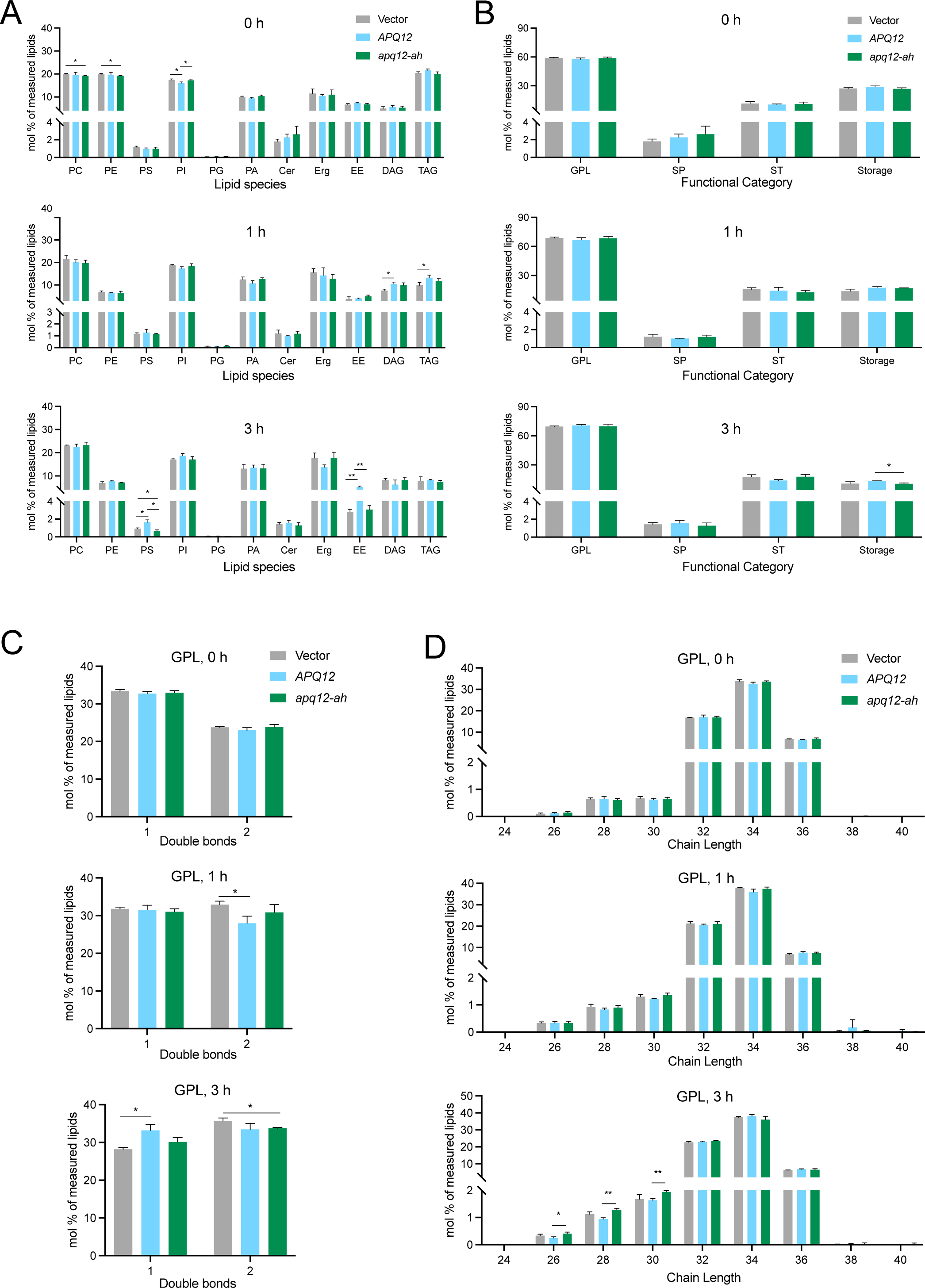
*APQ12* overexpression induces lipid metabolism flow from storage lipids towards membrane lipids. (A-D) Samples were prepared from cells incubated at 30°C and expressing PGal1-*APQ12- 6His* and PGal1-*Apq12-ah-6His* upon addition of galactose for indicated times, and lipids were extracted for subsequent analysis by nano-ESI-MS/MS. (A) Plots of lipid species in mol% of measured lipids. (B) Plots of functional categories of lipids; glycerophospholipids and glycerolipids (GPL), sphingolipids (SP), sterols (ST), and storage lipids. (C) Plots showing the number of double bonds in GPL. (D) Plots showing the chain lengths of the GPL. n = 3. Values are given as means + SD. t-test *p<0.05, **p<0.01.

**Figure S7.**
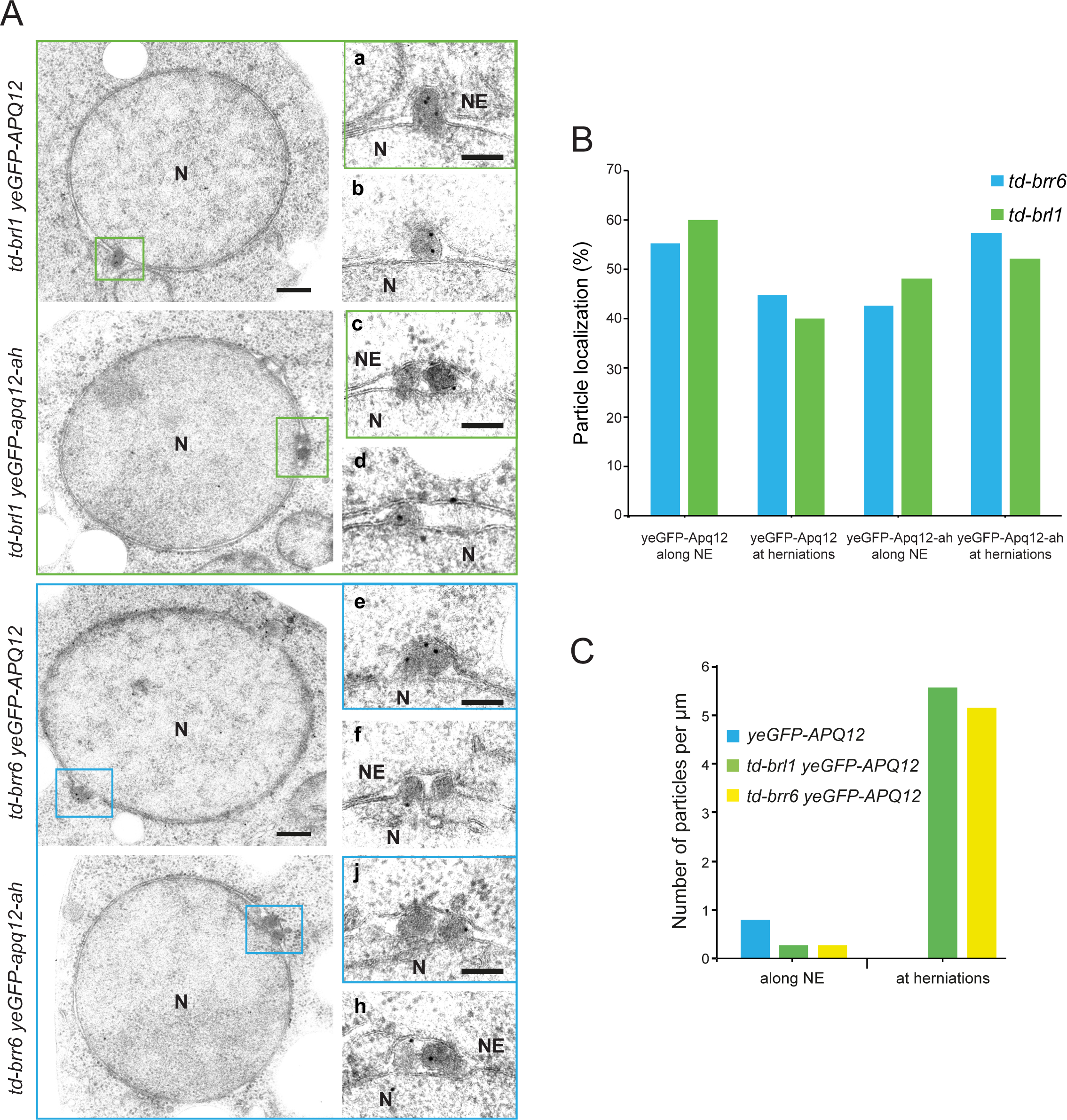
Localization of Apq12-yeGFP with herniations. (A) Localization of yeGFP-Apq12 and yeGFP-Apq12-ah was analyzed in *td-brl1* and *td-brr6* cells by immuno-EM as described in Fig. 8G. The 10 nm gold particles indicate localization of yeGFP-Apq12 and yeGFP-Apq12-ah. Scale bars: 200 nm, enlargements 100 nm. (B) Quantification of data from Fig. 8G and Fig. S7A. Gold particles reflecting localization of yeGFP-Apq12 and yeGFP-Apq12-ah at herniations were classified as indicated in the figure. Localization of 31 gold particles was analyzed for *td-brr6 APQ12-yeGFP*, 38 for *td-brl1 APQ12-yeGFP*, 37 for *td-brr6 apq12-ah-yeGFP* and 33 for *td-brl16 apq12-ah-yeGFP* cells. (C) Graph quantifying the number of particles, reflecting yeGFP-Apq12 and yeGFP-Apq12- ah, per µm of the NE. 24 gold particles were analyzed for *yeGFP-APq12*, 32 for *td-brl1 yeGFP-APQ12* and 39 for *td-brr6 yeGFP-APQ12* cells.

**Table S1:**
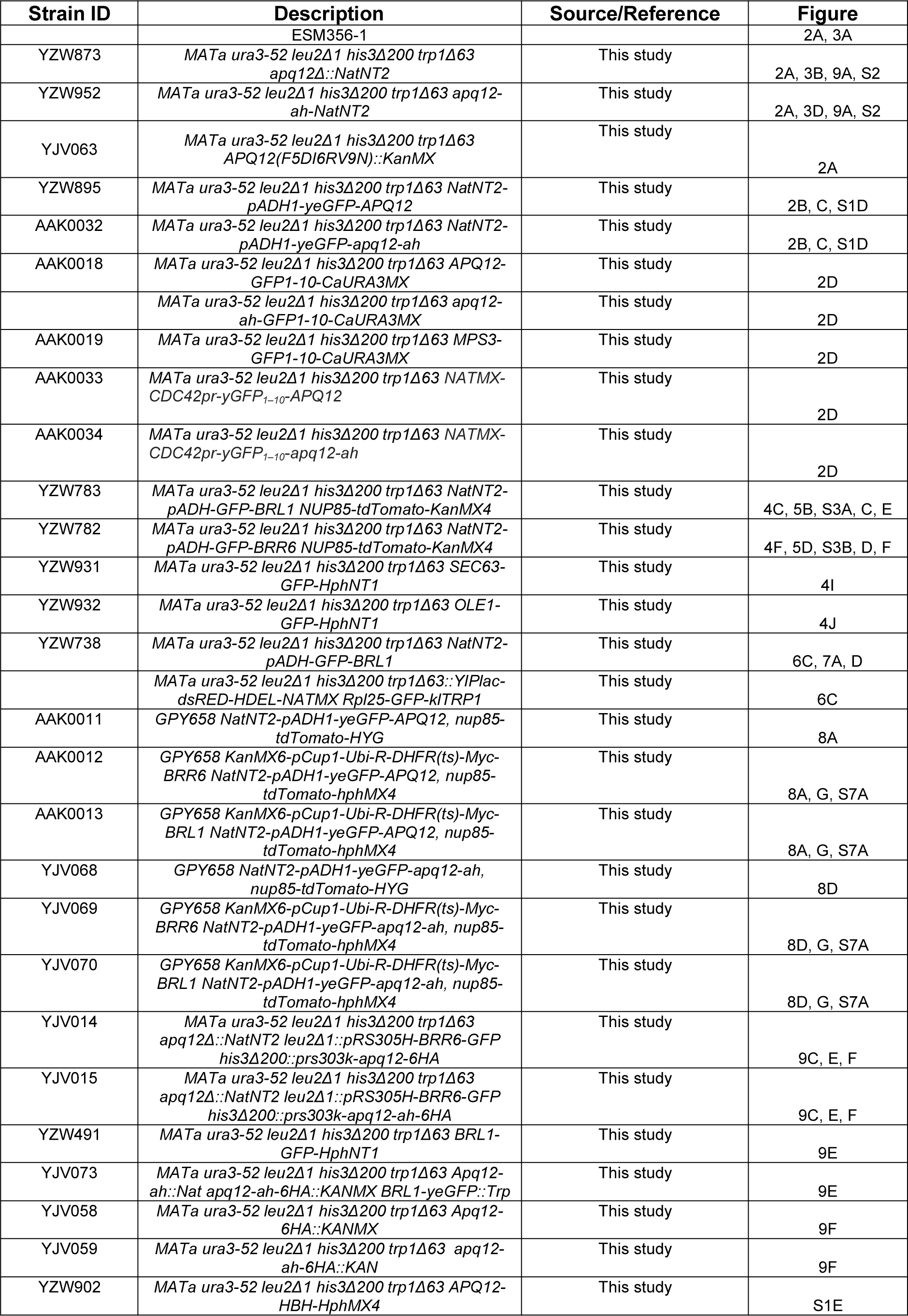
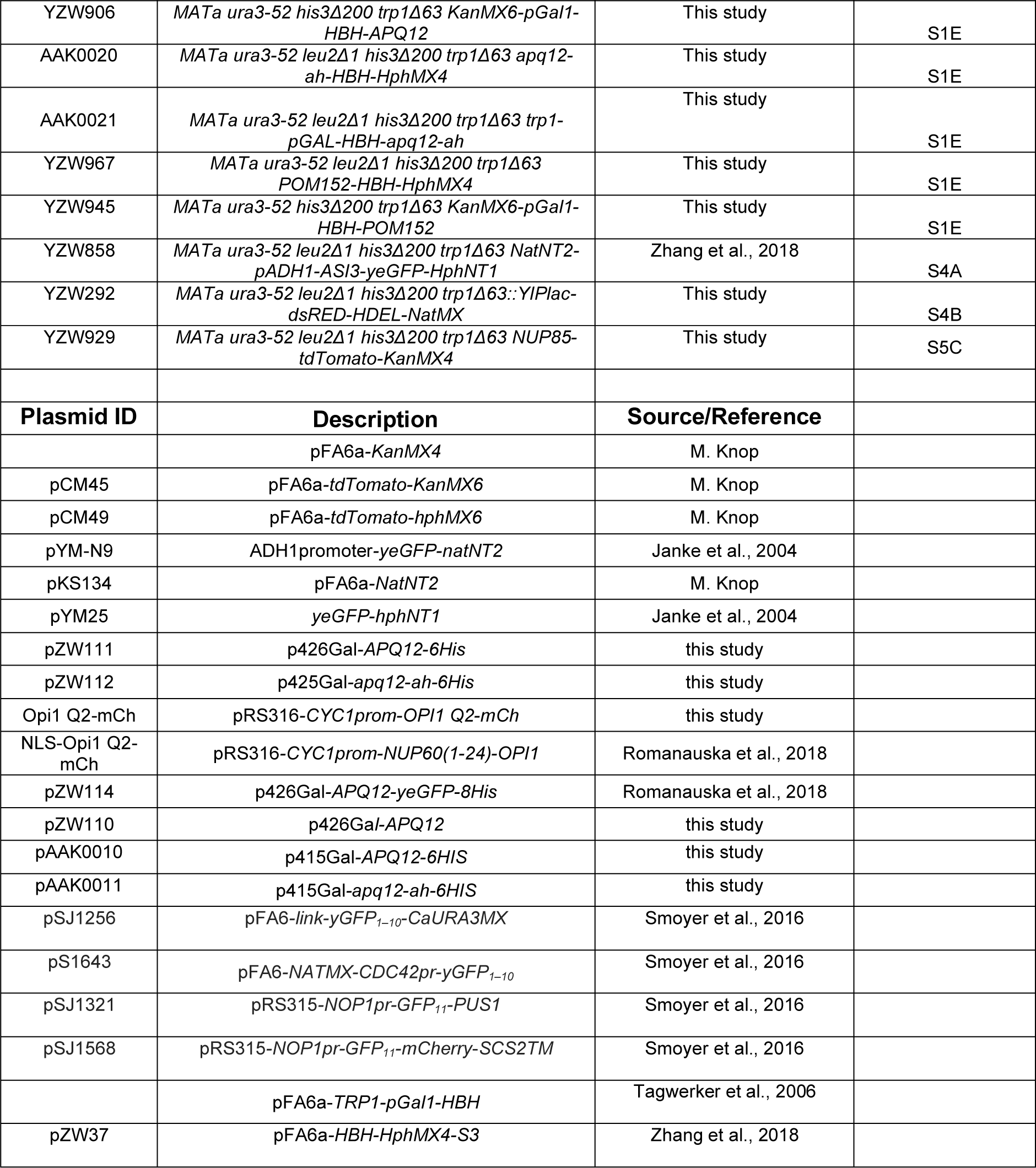
Yeast strains and plasmids used in this study.

## References

Aitchison JD, Blobel G, Rout MP (1995) Nup120p: A yeast nucleoporin required for NPC distribution and mRNA transport. Journal of Cell Biology 131: 1659–1675

Allegretti M, Zimmerli CE, Rantos V, Wilfling F, Ronchi P, Fung HKH, Lee CW, Hagen W, Turonova B, Karius K, Bormel M, Zhang XJ, Muller CW, Schwab Y, Mahamid J, Pfander B, Kosinski J, Beck M (2020) In-cell architecture of the nuclear pore and snapshots of its turnover. Nature 586: 796-+

Beck M, Hurt E (2017) The nuclear pore complex: understanding its function through structural insight. Nat Rev Mol Cell Biol 18: 73–89

Borah S, Thaller JD, Hakhverdyan Z, Rodriguez CE, Isenhour WA, Rout MR, King CM, Lusk CP (2021) Heh2/Man1 may be an evolutionarily conserved sensor of NPC assembly state. Mol Biol Cell 32

Carpenter AE, Jones TR, Lamprecht MR, Clarke C, Kang IH, Friman O, Guertin DA, Chang JH, Lindquist RA, Moffat J, Golland P, Sabatini DM (2006) CellProfiler: image analysis software for identifying and quantifying cell phenotypes. Genome Biology 7

de Bruyn Kops A, Guthrie C (2001) An essential nuclear envelope integral membrane protein, Brr6p, required for nuclear transport. EMBO J 20: 4183–93

Doucet CM, Talamas JA, Hetzer MW (2010) Cell cycle-dependent differences in nuclear pore complex assembly in metazoa. Cell 141: 1030–41

Eisenberg D, Weiss RM, Terwilliger TC (1982) The Helical Hydrophobic Moment - a Measure of the Amphiphilicity of a Helix. Nature 299: 371–374

Ejsing CS, Sampaio JL, Surendranath V, Duchoslav E, Ekroos K, Klemm RW, Simons K, Shevchenko A (2009) Global analysis of the yeast lipidome by quantitative shotgun mass spectrometry. Proceedings of the National Academy of Sciences of the United States of America 106: 2136–2141

Feldheim D, Rothblatt J, Schekman R (1992) Topology and functional domains of Sec63p, an endoplasmic reticulum membrane protein required for secretory protein translocation. Mol Cell Biol 12: 3288–96

Giddings TH, Jr., O’Toole ET, Morphew M, Mastronarde DN, McIntosh JR, Winey M (2001) Using rapid freeze and freeze-substitution for the preparation of yeast cells for electron microscopy and three-dimensional analysis. Methods Cell Biol 67: 27–42

Gimenez-Andres M, Copic A, Antonny B (2018) The Many Faces of Amphipathic Helices. Biomolecules 8

Guthrie C, Fink GR (1991) Guide to yeast genetics and molecular biology. Meth Enzymol 194: 429–663

Hashemi HF, Goodman JM (2015) The life cycle of lipid droplets. Curr Opin Cell Biol 33: 119–24

Hodge CA, Choudhary V, Wolyniak MJ, Scarcelli JJ, Schneiter R, Cole CN (2010) Integral membrane proteins Brr6 and Apq12 link assembly of the nuclear pore complex to lipid homeostasis in the endoplasmic reticulum. J Cell Sci 123: 141–51

Hu JJ, Shibata Y, Voss C, Shemesh T, Li ZL, Coughlin M, Kozlov MM, Rapoport TA, Prinz WA (2008) Membrane proteins of the endoplasmic reticulum induce high- curvature tubules. Science 319: 1247–1250

Jang JH, Lee CS, Hwang D, Ryu SH (2012) Understanding of the roles of phospholipase D and phosphatidic acid through their binding partners. Progress in Lipid Research 51: 71–81

Janke C, Magiera MM, Rathfelder N, Taxis C, Reber S, Maekawa H, Moreno- Borchart A, Doenges G, Schwob E, Schiebel E, Knop M (2004) A versatile toolbox for PCR-based tagging of yeast genes: new fluorescent proteins, more markers and promoter substitution cassettes. Yeast 21: 947–962

Khmelinskii A, Blaszczak E, Pantazopoulou M, Fischer B, Omnus DJ, Le Dez G, Brossard A, Gunnarsson A, Barry JD, Meurer M, Kirrmaier D, Boone C, Huber W, Rabut G, Ljungdahl PO, Knop M (2014) Protein quality control at the inner nuclear membrane. Nature 516: 410–3

Knop M, Siegers K, Pereira G, Zachariae W, Winsor B, Nasmyth K, Schiebel E (1999) Epitope tagging of yeast genes using a PCR-based strategy: More tags and improved practical routines. Yeast 15: 963–972

Lone MA, Atkinson AE, Hodge CA, Cottier S, Martinez-Montanes F, Maithel S, Mene- Saffrane L, Cole CN, Schneiter R (2015) Yeast Integral Membrane Proteins Apq12, Brl1, and Brr6 Form a Complex Important for Regulation of Membrane Homeostasis and Nuclear Pore Complex Biogenesis. Eukaryot Cell 14: 1217–27

Malsam J, Parisotto D, Bharat TAM, Scheutzow A, Krause JM, Briggs JAG, Sollner TH (2012) Complexin arrests a pool of docked vesicles for fast Ca2+-dependent release. Embo Journal 31: 3270–3281

Meijer M, Dorr B, Lammertse HCA, Blithikioti C, van Weering JRT, Toonen RFG, Sollner TH, Verhage M (2018) Tyrosine phosphorylation of Munc18-1 inhibits synaptic transmission by preventing SNARE assembly. Embo Journal 37: 300–320

Murphy R, Watkins JL, Wente SR (1996) GLE2, a Saccharomyces cerevisiae homologue of the Schizosaccharomyces pombe export factor RAE1, is required for nuclear pore complex structure and function. Molecular Biology of the Cell 7: 1921–1937

Onischenko E, Noor E, Fischer JS, Gillet L, Wojtynek M, Vallotton P, Weis K (2020) Maturation Kinetics of a Multiprotein Complex Revealed by Metabolic Labeling. Cell 183: 1785-+

Onischenko E, Tang JH, Andersen KR, Knockenhauer KE, Vallotton P, Derrer CP, Kralt A, Mugler CF, Chan LY, Schwartz TU, Weis K (2017) Natively Unfolded FG Repeats Stabilize the Structure of the Nuclear Pore Complex. Cell 171: 904–917 e19

Otsuka S, Bui KH, Schorb M, Hossain MJ, Politi AZ, Koch B, Eltsov M, Beck M, Ellenberg J (2016) Nuclear pore assembly proceeds by an inside-out extrusion of the nuclear envelope. Elife 5

Otsuka S, Ellenberg J (2018) Mechanisms of nuclear pore complex assembly - two different ways of building one molecular machine. FEBS Lett 592: 475–488

Otsuka S, Steyer AM, Schorb M, Heriche JK, Hossain MJ, Sethi S, Kueblbeck M, Schwab Y, Beck M, Ellenberg J (2018) Postmitotic nuclear pore assembly proceeds by radial dilation of small membrane openings. Nat Struct Mol Biol 25: 21–28

Ozbalci B, Boyaci IH, Topcu A, Kadilar C, Tamer U (2013) Rapid analysis of sugars in honey by processing Raman spectrum using chemometric methods and artificial neural networks. Food Chem 136: 1444–1452

Rampello AJ, Laudermilch E, Vishnoi N, Prophet SM, Shao L, Zhao C, Lusk CP, Schlieker C (2020) Torsin ATPase deficiency leads to defects in nuclear pore biogenesis and sequestration of MLF2. J Cell Biol 219

Romanauska A, Kohler A (2018) The Inner Nuclear Membrane Is a Metabolically Active Territory that Generates Nuclear Lipid Droplets. Cell 174: 700–715 e18

Saitoh YH, Ogawa K, Nishimoto T (2005) Brl1p - a novel nuclear envelope protein required for nuclear transport. Traffic 6: 502–517

Sapay N, Guermeur Y, Deleage G (2006) Prediction of amphipathic in-plane membrane anchors in monotopic proteins using a SVM classifier. BMC Bioinformatics 7: 255

Scarcelli JJ, Hodge CA, Cole CN (2007) The yeast integral membrane protein Apq12 potentially links membrane dynamics to assembly of nuclear pore complexes. J Cell Biol 178: 799–812

Smoyer CJ, Katta SS, Gardner JM, Stoltz L, McCroskey S, Bradford WD, McClain M, Smith SE, Slaughter BD, Unruh JR, Jaspersen SL (2016) Analysis of membrane proteins localizing to the inner nuclear envelope in living cells. Journal of Cell Biology 215: 575–590

Sorger D, Daum G (2003) Triacylglycerol biosynthesis in yeast. Applied Microbiology and Biotechnology 61: 289–299

Souquet B, Freed E, Berto A, Andric V, Auduge N, Reina-San-Martin B, Lacy E, Doye V (2018) Nup133 Is Required for Proper Nuclear Pore Basket Assembly and Dynamics in Embryonic Stem Cells. Cell Reports 23: 2443–2454

Tagwerker C, Zhang H, Wang X, Larsen LS, Lathrop RH, Hatfield GW, Auer B, Huang L, Kaiser P (2006) HB tag modules for PCR-based gene tagging and tandem affinity purification in *Saccharomyces cerevisiae*. Yeast 23: 623–32

Talamas JA, Hetzer MW (2011) POM121 and Sun1 play a role in early steps of interphase NPC assembly. Journal of Cell Biology 194: 27–37

Tamm T, Grallert A, Grossman EP, Alvarez-Tabares I, Stevens FE, Hagan IM (2011) Brr6 drives the *Schizosaccharomyces pombe* spindle pole body nuclear envelope insertion/extrusion cycle. J Cell Biol 195: 467–84

Tanguy E, Kassas N, Vitale N (2018) Protein-Phospholipid Interaction Motifs: A Focus on Phosphatidic Acid. Biomolecules 8

Tcheperegine SE, Marelli M, Wozniak RW (1999) Topology and functional domains of the yeast pore membrane protein Pom152p. J Biol Chem 274: 5252–8

Thaller DJ, Allegretti M, Borah S, Ronchi P, Beck M, Lusk CP (2019) An ESCRT- LEM protein surveillance system is poised to directly monitor the nuclear envelope and nuclear transport system. Elife 8

Thaller DJ, Tong DQ, Marklew CJ, Ader NR, Mannino PJ, Borah S, King MC, Ciani B, Lusk CP (2021) Direct binding of ESCRT protein Chm7 to phosphatidic acid-rich membranes at nuclear envelope herniations. Journal of Cell Biology 220

Thiam AR, Farese RV, Jr., Walther TC (2013) The biophysics and cell biology of lipid droplets. Nat Rev Mol Cell Biol 14: 775–86

Ungricht R, Kutay U (2017) Mechanisms and functions of nuclear envelope remodelling. Nat Rev Mol Cell Biol 18: 229–245

van Leeuwen J, Pons C, Tan GH, Wang JSZ, Hou J, Weile J, Gebbia M, Liang W, Shuteriqi E, Li ZJ, Lopes M, Usaj M, Lopes AD, van Lieshout N, Myers CL, Roth FP, Aloy P, Andrews BJ, Boone C (2020) Systematic analysis of bypass suppression of essential genes. Molecular Systems Biology 16

Wang N, Clark LD, Gao Y, Kozlov MM, Shemesh T, Rapoport TA (2021) Mechanism of membrane-curvature generation by ER-tubule shaping proteins. Nat Commun 12

Weber T, Zemelman BV, McNew JA, Westermann B, Gmachl M, Parlati F, Sollner TH, Rothman JE (1998) SNARE dependent membrane fusion. Mol Biol Cell 9: 331a- 331a

Webster BM, Colombi P, Jager J, Lusk CP (2014) Surveillance of nuclear pore complex assembly by ESCRT-III/Vps4. Cell 159: 388–401

Wente SR, Blobel G (1993) A temperature-sensitive NUP116 null mutant forms a nuclear envelope seal over the yeast nuclear pore complex thereby blocking nucleocytoplasmic traffic. J Cell Biol 123: 275–84

Winey M, Hoyt MA, Chan C, Goetsch L, Botstein D, Byers B (1993) NDC1: a nuclear periphery component required for yeast spindle pole body duplication. J Cell Biol 122: 743–751

Winey M, Yarar D, Giddings TH, Jr., Mastronarde DN (1997) Nuclear pore complex number and distribution throughout the Saccharomyces cerevisiae cell cycle by three-dimensional reconstruction from electron micrographs of nuclear envelopes. Mol Biol Cell 8: 2119–32

Zhang W, Neuner A, Rüthnick D, Sachsenheimer T, Lüchtenborg C, Brügger B, Schiebel E (2018) Brr6 and Brl1 locate to nuclear pore complex assembly sites to promote their biogenesis. J Cell Biol 217: 877–894

Zhendre V, Grelard A, Garnier-LHomme M, Buchoux S, Larijani B, Dufourc EJ (2011) Key Role of Polyphosphoinositides in Dynamics of Fusogenic Nuclear Membrane Vesicles. Plos One 6

Zhukovsky MA, Filograna A, Luini A, Corda D, Valente C (2019) Phosphatidic acid in membrane rearrangements. Febs Letters 593: 2428–2451

